# A generic reference defined by consensus peaks for scATAC-seq data analysis

**DOI:** 10.1101/2023.05.30.542889

**Authors:** Qiuchen Meng, Xinze Wu, Wenchang Chen, Yubo Zhao, Chen Li, Zheng Wei, Jiaqi Li, Xi Xi, Sijie Chen, Catherine Zhang, Shengquan Chen, Jiaqi Li, Xiaowo Wang, Rui Jiang, Lei Wei, Xuegong Zhang

## Abstract

The rapid advancement of transposase-accessible chromatin using sequencing (ATAC-seq) technology, particularly with the emergence of single-cell ATAC-seq (scATAC-seq), has accelerated the studies of gene regulation. However, the absence of a generic feature reference for ATAC-seq data limits single-cell analyses and hinders the development of comprehensive cell atlases. To address this, we constructed a generic chromatin accessibility reference by aggregating peaks from 624 high-quality bulk ATAC-seq datasets, defining more than 1 million consensus peaks (cPeaks). Leveraging a deep neural network model, we expanded cPeaks to include previously unobserved regions, enhancing their coverage across diverse tissues and cell types. cPeaks exhibit consistent shapes across tissue types, sequencing technologies, and peak-calling methods, indicating that they represent inherent genomic features. Compared to existing feature defining methods and references, cPeaks show superior performance in scATAC-seq analyses, improving cell annotation and rare cell type identification. Additionally, cPeaks provide insights into chromatin dynamics during cellular differentiation and tumor progression. cPeaks can serve as a robust reference for chromatin accessibility studies to promote cross-dataset consistency and accelerate biological discoveries.

## Introduction

Chromatin accessibility refers to the availability of chromatinized DNA to directly interact with other macromolecules^1^. The accessible regions^2,3^, also known as open regions^1,2^, often associate with regulatory elements such as promoters, enhancers, insulators, and transcription factor (TF) binding sites (TFBSs)^4,5^. Studying accessible regions is crucial for deciphering gene regulations^2,6–8^ across various biological processes, including cell differentiation^9,10^, organ development^11,12^, and disease manifestation^13–16^. The advancements in high-throughput sequencing technologies^3^, particularly Assay for Transposase-Accessible Chromatin using sequencing (ATAC- seq)^17,18^, have revolutionized our ability to measure chromatin accessibility on a genome-wide scale by offering more efficient and simplified approaches. Especially, the emergence of single-cell ATAC-seq (scATAC-seq)^19–24^ now allows for the exploration of chromatin accessibility at single-cell resolution, complementing single-cell RNA sequencing (scRNA-seq) for investigating the molecular properties of cells of different types or states and for building the biomolecular atlases of cells^5,16,25,26^.

Unlike RNA-seq, which benefits from established reference transcriptomes^27^, ATAC-seq lacks a standardized reference for localizing and quantifying chromatin accessibility across the genome. Traditional approaches in bulk ATAC-seq involve creating sample-specific feature sets, using peak-calling algorithms such as MACS2^28^ and Cell Ranger ATAC^29^ to identify regions with significantly enriched read signals as accessible regions or peaks^5,25,30^.

When integrating data from different samples, current practices usually first call peaks as features in each dataset, and then integrate these features with the peak merging^31,32^ or iterative removal^5,16^ approach. The merging approach combines overlapping peaks into broader regions, potentially merging adjacent peaks inaccurately. The iterative removal approach selects the most significant peak from overlapping areas, risking the loss of valuable information.

The advent of scATAC-seq provides a single-cell resolution of chromatin accessibility information and brings substantial increases in the scale of the data, often comprising tens to hundreds of thousands of cells^5,26,30^. However, peak-calling at the single-cell level is impractical, necessitating the merging of multiple cells into pseudo-bulk samples to define accessible regions^5,25,26,30,33,34^. This results in the loss of single-cell resolution and potentially obscures unique chromatin features of rare cell types. To improve this, iterative techniques for clustering^5^ and optimized latent semantic indexing (LSI)^35^ have been developed to improve the identification of accessible regions, especially in rare cell types. Nevertheless, these techniques still involve multiple rounds of feature integration, which increases with scATAC-seq data scales, leading to labor-intensive processes and aggrevated information loss, especially in atlas-scale analyses.

These challenges highlight the critical need for a generic reference of accessible regions to standardize analyses across the increasing volume of sequencing data, and to enhance the consistency and accuracy of research. Valentina et al. have shown that using predefined sets of genomic regions can facilitate the mapping of scATAC-seq data^36^.

This underscores the potential transformative impact of a comprehensive reference that encompasses all or at least the major potential accessible regions shared across datasets on scATAC-seq data analysis. Currently, several pre- defined sets of accessible regions have been proposed to identify and characterize possible gene regulatory elements^5,37,38^. For example, the consensus DNase I hypersensitive site (cDHS)^37^ offers a standard index of DNase I hypersensitive sites identified through DNaseLI hypersensitive seq (DNase-seq) data, and the *cis*-element ATLAS (CATLAS)^5^ was introduced as a repository of candidate *cis*-regulatory elements (cCREs) among the genome. cDHS is specifically based on DNase-seq technology, and CATLAS cCREs do not retain information on the length of accessible regions. Consequently, these resources are often employed for quality control in data analysis rather than as comprehensive references for data analysis^3^.

To construct a robust chromatin accessibility reference, we collected 624 high-quality public bulk ATAC-seq datasets, covering diverse tissue types. These datasets were used to defined approximately 1.4 million observed consensus peaks (cPeaks) as a chromatin accessibility reference. To evaluate the generalizability of cPeaks as a universal feature set for scRNA-seq data, we validated them using an independent set of 231 scATAC-seq datasets. We found consistent patterns in cPeak shapes across tissue types, sequencing technologies, and peak-calling methods, suggesting that cPeaks represent inherent genomic features. To address the limitation of unseen tissue-specific accessible regions in existing data, we developed a deep neural network model to predict new cPeaks, adding ∼0.28 million predicted regions to the reference. We used an additional independent set of 789 bulk ATAC-seq datasets to validate the reliability of predicted cPeaks. The combined observed and predicted cPeaks serve as a standardized set of chromatin accessibility reference, enhancing cross-dataset consistency in ATAC-seq analyses. Compared to existing feature defining methods and references, cPeaks significantly improved scATAC-seq cell annotation and rare cell type identification. Additionally, cPeaks facilitated the exploration of chromatin dynamics in complex biological states such as cellular differentiation and tumor progression. We provided comprehensive resources and bioinformatics tools to integrate cPeaks into multiple analysis pipelines to enable their effective use in all types of chromatin accessibility studies.

## Results

### Defining cPeaks by aggregating ATAC-seq data

We used bulk ATAC-seq data to establish a robust chromatin accessibility reference. We collected all available bulk ATAC-seq datasets derived from healthy adult samples from two widely used repositories, ENCODE^39^ and ATACdb^40^. After quality control, the collection resulted in a total of 624 high-quality datasets, covering over 40 human organs and all body systems (Methods, **Table S1**). Each dataset provided a set of pre-identified peaks, each marking an accessible region on the genome observed in respective dataset (**Fig. 1a**). Peaks were iteratively merged, one dataset at a time, with non-overlapping peaks directly added to the pool and overlapping peaks combined into broader regions (Methods). As more datasets were integrated, the total number of peaks and their genomic coverage initially increased but eventually tended to reach a plateau (**Fig. 1b**), suggesting that the total number of all possible accessible regions on the genome may be limited.

**Fig 1.**
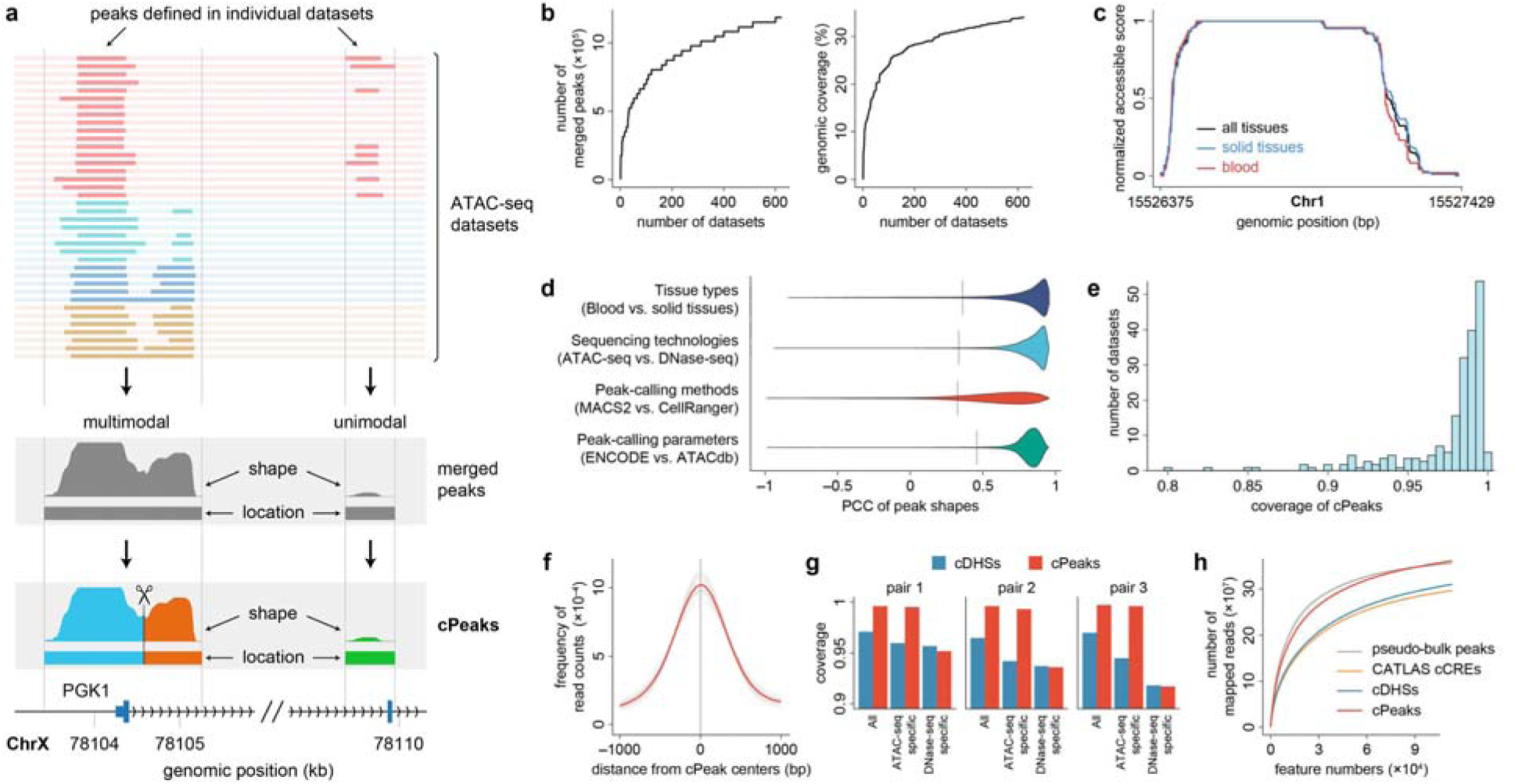
The computation diagram of cPeaks and their basic properties. (a) The schematic diagram of cPeak generation. A part of the genomic region near *PGK1* is shown as an example. In the upper panel, each row represents an ATAC-seq dataset, and each segment represents a pre-identified peak. Different colors indicate different cell types. The middle panel illustrates a multimodal merged peak (left) and a unimodal merged peak (right). In the bottom panel, the multimodal merged peak is clipped as two cPeaks, and the unimodal merged peak forms one cPeak. The relative positions of the cPeaks with regard to the transcription start site (TSS) and the 2nd exon of PGK1 are shown at the bottom. (b) Increase tendency of the number of merged peaks (left) and their genomic coverage (right) with the addition of datasets to be aggregated. (c) An example of the shapes of merged peaks on chromosome 1, obtained with data of only blood samples (red line), of only solid tissues (blue line), or of all tissues (black line). We normalized the accessible score such that the maximum score scaled to 1. (d) Correlations of peak shapes between different conditions. The grey line shows the average median PCC from 100 permutation tests by randomly shuffling merged peak pairs. (e) Histogram of the coverages of cPeaks in 231 scATAC-seq datasets. (f) Enrichment of sequencing reads around centers of cPeaks. All sequencing reads overlapped with ±1,000 bp genomic regions around cPeak centers are aggregated to form the distributions. Each grey line represents a dataset. The red line shows the average distribution of all datasets. (g) The coverage of cPeaks and cDHSs on different categories of accessible regions. Different blocks represent different pairs of ATAC-seq and DNase-seq data. “All” represents represents all accessible regions detected in the ATAC-seq data, while “ATAC- seq/DNase-seq specific” represents the unique accessible regions detected by each sequencing technology. (h) The change of the number of mapped reads by adding sorted features. The order of features is sorted from high to low according to the number of mapped reads.

After processing all datasets, we identified approximately 1.2 million merged peaks, covering about 30% of the whole genome. Each merged peak represents a genomic region that could be accessible in one or more cell types. For each position within a merged peak, we calculated the proportion of datasets in which the position was covered by peaks, defining this proportion as the “accessible score” for the position. The series of accessible scores along the 5’-to-3’ direction of a merged peak was summarized into a profile referred to as the “shape” of the peak (Methods).

We then investigated whether the shapes of a merged peak are consistent under varying conditions, including tissue types, sequencing technologies, and peak-calling methods. First, we examined whether tissue types influenced peak shapes. The datasets used to generate merged peaks were derived from two major tissue categories: about 51% from blood and others from solid tissues (e.g., lung, heart, brain). We used the same procedure to merge peaks for these two tissue types separately. Among the merged peaks, 87% and 76% were located at shared genomic regions between the blood and solid tissue datasets, respectively. We found the shared peaks exhibited similar shapes, as illustrated in **Fig. 1c**. We quantified shape similarity for these shared peaks using the Pearson correlation coefficient (PCC) between their shapes in blood and solid tissue data, obtaining a median PCC of 0.9 (**Fig. 1d**, Methods). Next, we examined whether sequencing technologies affected peak shapes by comparing merged peak shapes with those generated from DNase-seq data (cDHS)^37^. The results demonstrated high similarity in shapes at corresponding genomic regions (**Fig. 1d**, Methods). Lastly, we tested whether peak-calling methods and parameter variations influenced shapes by comparing peaks generated by CellRanger^29^ and MACS2^28^ different parameters (Methods).

The results showed a high consistency of shapes across different conditions (**Fig. 1d**). These findings highlight the consistent shapes of merged peaks, regardless of tissue type, sequencing technology, or peak-calling methodology. This suggests that merged peaks may represent an inherent genomic feature and can serve as a basis for generating a chromatin accessibility reference.

We found that merged peaks exhibited two distinct shapes: unimodal or multimodal (**Fig. 1a**, Methods). We found that multimodal merged peaks were composed of adjacent unimodal peaks and therefore we decomposed them into multiple unimodal peaks (**Fig. 1a**, Methods). These unimodal merged peaks were designated as the basic unit of the chromatin accessibility reference, termed observed consensus peaks (cPeaks). In this way, we obtained approximately 1.4 million observed cPeaks in total (an example provided in **Fig. S1a**). The majority of observed cPeaks were shorter than 2,000 bp, with a median length of 525 bp, closely aligning with previous findings that ATAC-seq peaks typically span around 500 bp^41^ (**Fig. S1b**).

We assessed the feasibility of cPeaks as a chromatin accessibility reference for processing scATAC-seq data. To ensure broad validataion, we collected 231 scATAC-seq datasets from published studies (**Table S2**), each containing peaks pre-identified using different peak-calling methods^28,29^. For each dataset, we calculated the proportion of the pre-identified peaks overlapping with observed cPeaks, referred to as cPeak coverage. Most datasets exhibited coverage above 0.9, with an average of 0.98 and a median of 0.99 (**Fig. 1e**). Observed cPeaks consistently maintained high coverage across different tissue types, pre-identified peak numbers, and peak-calling parameters without clear bias (**Fig. S1c**).

To further evaluate the enrichment of sequencing reads on observed cPeaks, we used the 27 scATAC-seq datasets from CATLAS^5^ to invest cPeaks’ performance across different organs. We found that the sequencing reads were predominantly enriched near the centers of observed cPeaks (**Fig. 1f**, Methods). On average, 68.8% of sequencing reads in these scATAC-seq data were mapped to observed cPeaks, surpassing the typical mapping rate of ∼65% reported in the technical reports of 10X Genomics^42,43^. These findings establish cPeaks as a reliable reference for capturing the signals in scATAC-seq data across diverse organs.

We compared the performance of cPeaks against other pre-defined accessible region sets, including cDHS and CATLAS cCREs. First, we assessed the differences in coverage between cPeaks and cDHS on accessible regions identified by ATAC-seq and DNase-seq. By analyzing three additional ATAC-seq and three DNase-seq data from the GM12878 cell line^44^, we found that compared to cDHS, cPeaks provided higher coverage of ATAC-seq-specific accessible regions while maintaining similar coverage for DNase-seq-unique regions (**Fig. 1g**, Methods). This demonstrates that cPeaks encompass a broader range of accessible regions detected by ATAC-seq compared to cDHS.

We then assessed the coverage of scATAC-seq sequencing reads on accessible regions defined by cPeaks, cDHS, and CATLAS cCREs using a hematopoietic differentiation dataset^9^. We identified peaks through MACS2^28^ peak calling results as the ground truth of accessible regions. For each pre-identified accessible region set, we mapped sequencing reads to the set and sorted accessible regions by the number of mapped reads. As shown in **Fig. 1h**, the ratio of aggregated mapped reads using cPeaks closely aligned the ground truth, significantly exceeding those obtained with CATLAS cCREs and cDHS. The results suggested that cPeaks may serve as a more comprehensive and appropriate reference for scATAC-seq data analysis.

### Expanding cPeaks through deep learning

cPeaks were derived from a finite set of ATAC-seq datasets. Despite the considerable sample size, concerns remain about their ability to handle unseen cell types with unique accessible regions. To address this limitation and enhance cPeaks as a universal chromatin accessibility reference, we employed a deep learning model to expand cPeaks. We hypothesized that cPeak positions are related to their DNA sequences, as the non-random distribution of cPeaks across the genome (**Fig. 1b**) and their consistent shapes, regardless of tissue type, sequencing technology, or peak- calling methodology, point to a sequence-dependent mechanism underlying their genomic positioning. We thus implemented a deep convolutional neural network (CNN) inspired by scBasset^45^ to classify genomic regions as cPeaks or non-cPeaks based on DNA sequence patterns (**Fig. 2a**). In this model, observed cPeaks served as positive samples, while randomly selected regions without cPeaks were used as negative controls (Methods). The model achieved a high accuracy with an AUC of 94.72% on test data (**Fig. 2b**), highlighting a strong relationship between cPeak locations and their DNA sequence signatures. These findings reinforce the hypothesis that cPeaks are an inherent genomic feature and demonstrate the feasibility of predicting unobserved cPeaks using DNA sequence information.

**Fig. 2.**
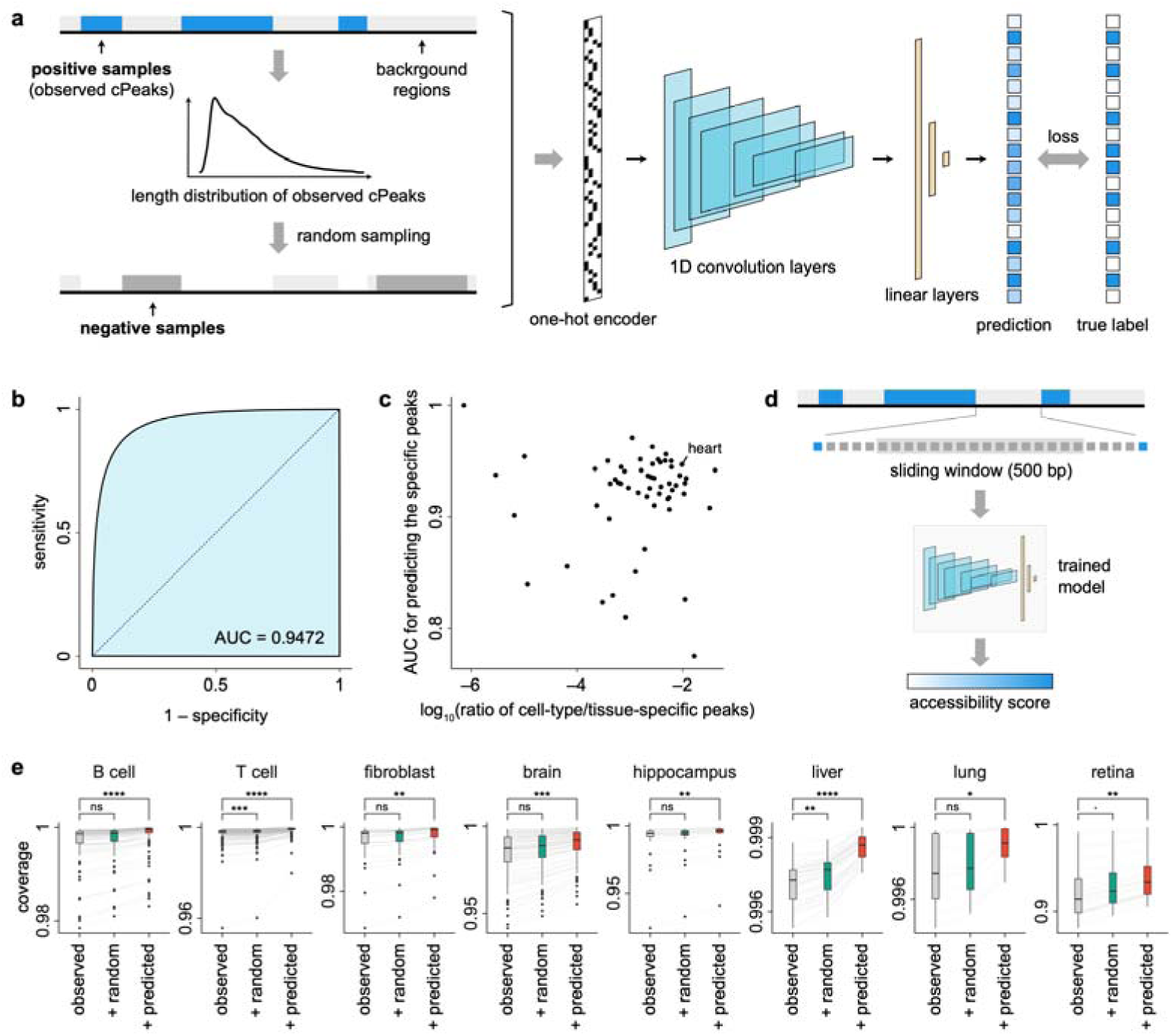
Extending cPeaks with deep learning. (a) The one-dimensional CNN model for predicting whether a genomic region is a cPeak. We considered cPeaks as positive samples and followed their length distribution to extra negative samples from background regions. (b) The ROC curve of the deep CNN model on test data. (c) The AUC for distinguishing excluded cell-type/tissue-specific cPeaks from an equal number of negative samples after training the CNN model without the corresponding cell-type/tissue-specific cPeaks. (d) Predicting cPeaks by the CNN model. We scanned all background regions by a window of 500 bp with a stride of 250 bp. Each region was fed into the trained model to get a predicted accessibility score. (e) Coverages of different ATAC-seq data by three sets of cPeaks: the observed cPeaks based on the initial data, the observed cPeaks plus a random set of regions from the genomic background, and the observed cPeaks plus the predicted cPeaks. Each grey line links the same dataset analyzed by different cPeak sets. The significant levels are calculated by the paired Wilcoxon signed-rank test (*p*- values denoted as ****, < 0.0001; ***, < 0.001; **, < 0.01; *, < 0.05; ·, < 0.1; ns, ≥ 0.1).

We evaluated the ability of the CNN model to predict missing cPeaks in cell types or tissues not represented during chromatin accessibility reference construction. Each cPeak is observed as accessible in one or more specific cell types or tissues, with cell-type/tissue-specific cPeaks defined as those unique to a single cell type or tissue. To simulate the absence of particular cell types or tissues during reference construction, we excluded their cell-type/tissue-specific cPeaks from training data while retaining the remaining cPeaks as positive samples. Negative samples were randomly selected from genomic regions without cPeaks, following the same procedure illustrated in **Fig. 2a**. The CNN model was then trained from scratch using these revised datasets (Methods). As shown in **Fig. 2c**, the model consistently achieved high prediction scores for distinguishing excluded cell-type/tissue-specific cPeaks from an equal number of negative samples, achieving a mean AUC of 91.67% across all evaluations. For instance, heart-specific cPeaks, which represent 0.97% of all cPeaks, were effectively identified with an AUC of 94.70% after training the model on the remaining 99.03% of cPeaks. These results indicate that the CNN model can effectively predict missing cPeaks for previously unseen tissues or cell types.

To expand the cPeak set, we used the trained CNN model to scan the entire human genome, applying a sliding window of 500 bp across all background regions. This approach assigned an accessibility score to each genomic region, representing its probability of being a cPeak (**Fig. 2d**, Methods). Regions were ranked by accessibility scores, with those scoring highest designated as predicted cPeaks, resulting in a total of approximately 0.28 million predicted cPeaks (Methods).

To evaluate the performance of predicted cPeaks, we collected an independent set of bulk ATAC-seq datasets from ChIP-Atlas^44^. After quality control, we obtained 789 datasets covering eight cell types or tissues: B cells, T cells, fibroblasts, brain, hippocampus, liver, lung, and retina (**Table S2**). These datasets were chosen to include both tissue types represented in the initial cPeak construction and a previously unseen tissue, retina. Importantly, none of these datasets were used in the initial construction process, ensuring an unbiased evaluation of predicted part. We also randomly generated a set of background regions with lengths and numbers matching the predicted cPeaks, referred to as random regions, to serve as random controls for comparison.

We evaluated the coverage of observed cPeaks alone, observed cPeaks plus random regions, and observed cPeaks plus predicted cPeaks across the 789 datasets. We found that observed cPeaks already covered more than 99.21% of the reported peaks for most datasets. For retina, the mean coverage of observed cPeaks was slightly lower at 92.24%. Adding random regions to observed cPeaks did not significantly improve coverage, whereas incorporating predicted cPeaks significantly increased coverage across most cell types and tissues (**Fig. 2e**). In previously observed tissues or cell types, datasets with initially lower observed coverage exhibited the greatest improvement, reducing coverage variance among samples after the inclusion of predicted cPeaks. For the unseen tissue, retina, the model demonstrated the most notable enhancement, with mean coverage increasing from 92.23% to 93.95% upon adding predicted cPeaks. These results indicate that the inclusion of predicted cPeaks enhances the ability of cPeaks to encompass accessible regions across diverse datasets, particularly benefiting cell types or tissues not initially represented in cPeak construction.

### Association between cPeaks with known regulatory elements and cellular functions

We explored the associations between cPeaks and known regulatory elements and biological functions. By comparing observed cPeaks with genomic annotations, we found that observed cPeaks are enriched near transcriptional start sites (TSSs) and transcription end sites (TESs) (**Fig. 3a**, Methods). Approximately 13% of observed cPeaks are localized at promoter regions (±2,000 bp around TSS), while the remaining cPeaks primarily distributed at exons or distal intergenic regions (**Fig. 3b**, Methods). We further collected multiple types of epigenetic annotations, including histone modification and CTCF ChIP-seq data (**Table S3**). Using these annotations, we found that most observed cPeaks contain or overlap with one or more regulatory elements, such as enhancers, promoters (both known and predicted), untranslated regions (UTRs), or CTCF binding sites (CTCF BSs) (**Fig. 3c**, Methods).

**Fig. 3.**
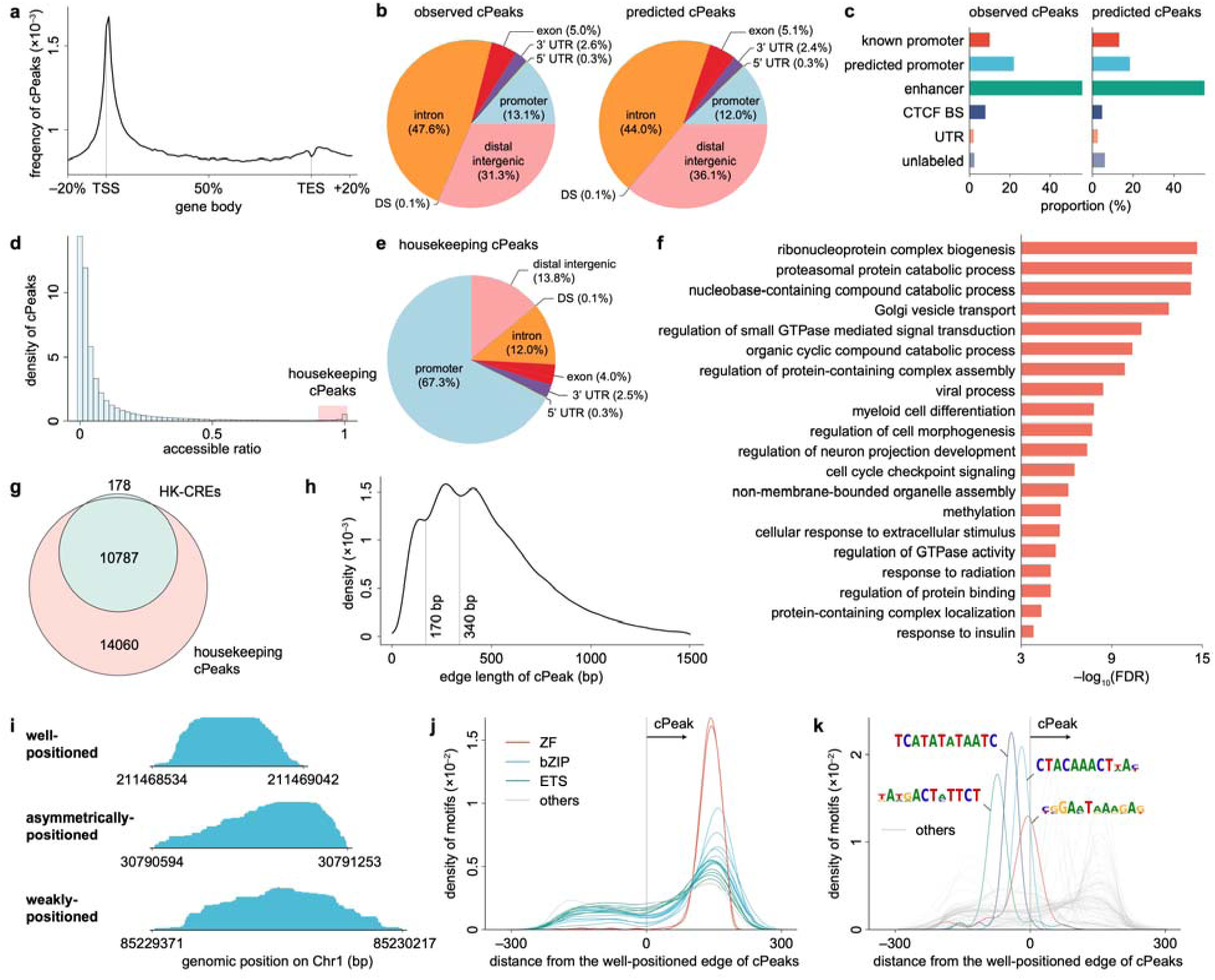
Properties of cPeaks. (a) The enrichment of cPeaks around gene bodies. The x-axis reflects the percentage of total length of gene body regions. The y-axis represents the frequency of observed cPeaks on each site. (b) Genomic locations of observed and predicted cPeaks. DS: downstream (≤ 300 bp downstream of the gene end). (c) The inferred regulatory functions of observed and predicted cPeaks. Note that a cPeak may have multiple inferred functions. CTCF BS: CTCF binding site. (d) The distribution of accessible ratios (proportions of tissues accessible at the cPeak location) of the observed cPeaks. We call the cPeaks with more than 0.9 accessible ratios (marked by the red box) as housekeeping cPeaks. (e) Genomic locations of housekeeping cPeaks. (f) Top 20 pathways enriched in genes related to housekeeping cPeaks. The 20 pathways with the smallest false discovery rates (FDRs) are shown. (g) Venn plot of housekeeping cPeaks and HK-CREs. (h) The distribution of the distribution of edge lengths of cPeaks. (i) Examples showing the three typical shape patterns of cPeaks. (j) Positions of motifs of the top 20 well- positioned-associated TFs. The x-axis is ±300 bp around the edges of cPeaks, from closed regions to accessible regions. Each line represents a motif, and each color represents a DNA-binding domain type. (k) Positions of the enriched *de novo* motifs around the well-positioned edges of cPeaks. The x-axis is ±300 bp around the edges of cPeaks, from closed regions to accessible regions. Each line represents a motif. We colored top 4 motifs on the closed regions around cPeaks.

Further genomic annotations and regulatory role inferences for predicted cPeaks revealed properties similar to those of observed cPeaks (**Fig. 3b-c**).

For each cPeak, we defined the maximum value of its shape as its “accessible ratio” to reflect the proportion of datasets where the cPeak is accessible. We defined a cPeak with an accessible ratio > 90% as a “housekeeping” cPeak (**Fig. 3d**). Among the observed cPeaks, 24,538 (1.8%) were identified as housekeeping cPeaks. The median length of housekeeping cPeaks was 1,441 bp, with most located at promoter regions (**Fig. 3e**). These promoter regions overlap with 77.4% of the promoters of housekeeping genes^46^ (Fisher’s exact test *p*-value < 1×10^−5^, Methods). Besides, the proportion of housekeeping cPeaks and the proportion of housekeeping genes across chromosomes were strongly correlated (**Fig. S2a**). Gene Ontology (GO) enrichment analysis of genes linked to housekeeping cPeaks revealed significant enrichment in essential biological pathways (Methods), including ribonucleoprotein complex biogenesis, cell cycle checkpoint signaling, and methylation (**Fig. 3f**). These findings align with those of a recently published study^47^ which proposed a similar concept Housekeeping CREs (HK-CREs) and reported that most HK-CREs are located in core promoters and associated with housekeeping gene-related biological processes. We found that housekeeping cPeaks encompass 98.4% of HK-CREs, underscoring their functional relevance (**Fig. 3g**).

We further examined the relationship between cPeak shapes and gene regulation. We defined the genomic region corresponding to the maximum value of each cPeak shape as its core accessible region. The distance from the endpoints of cPeak to the nearest boundary of the core accessible region is defined as the edge length. This metric reflects the consistency of accessible region boundaries across different cells. Smaller edge lengths indicated more consistent boundaries, suggesting the presence of well-positioned flanking nucleosomes, while larger edge lengths implied weakly-positioned flanking nucleosomes^1,48,49^. To ensure reliability, we filtered cPeaks with an accessible ratio greater than 0.3. As shown in **Fig. 3h**, the distribution of edge lengths exhibited three clusters with boundaries at 170 bp and 340 bp, approximately corresponding to the lengths of DNA wrapped around one and two nucleosomes, respectively. Edges shorter than 170 bp were categorized as well-positioned, while longer edges were classified as weakly-positioned. Based on edge type, cPeaks were further divided into three patterns: “well-positioned,” “asymmetrically-positioned,” and “weakly-positioned” (**Fig. 3i**).

We focused on well-positioned edges as they represent more precisely regulated regions. Two potential biological mechanisms were hypothesized to underlie the occurrence of well-positioned edges: the presence of specific TFBSs at the inner boundary and well-positioned flanking nucleosomes outside the boundary. To investigate the first hypothesis, we conducted motif enrichment analysis around well-positioned edges to identify well-positioned- associated TFs. These identified TFs included CTCF, BORIS, Fosl2, Fra2, Fli1, and ETS1 (**Fig. S2b**, Methods). The top 20 well-positioned-associated TFs shared common DNA-binding domains, including the basic leucine zipper (bZIP), Zinc finger (Zf), and erythroblast transformation specific (ETS) domains (**Fig. 3j**). Beyond the top 20, all well-positioned-associated TFs were also enriched for additional domains, such as homeobox and basic helix-loop-helix (bHLH) (**Fig. S2c**). The GO enrichment analysis result showed that these TFs were related to essential biological processes, including the regulation of transcription, cell development, and cell differentiation (**Fig. S2d**, Methods). We assumed that these well-positioned-associated TFs with these domains may possess stronger nucleosome remodeling capabilities. Besides, we found pioneer transcription factors (PTFs)^50–52^ were enriched in these TFs with a *p*-value of 0.016 (Fisher’s exact test, Methods). For the second hypothesis, we performed *de novo* motif enrichment analysis for the sequence of well-positioned edges, identifying sequence motifs associated with well-positioned flanking nucleosome positioning around cPeak boundaries (**Fig. 3k**, Methods). Together, well-positioned edges likely arise from the interplay of specific TFBSs and well-positioned flanking nucleosomes, highlighting their role in precise genomic regulation.

Most cPeaks are accessible in only a limited number of tissues or cell types (**Fig. 3d**), leaving 92.62% of cPeaks with unlabeled shapes due to low accessible ratios. To address this, we developed a neural network, a modified version of the CNN model used for cPeak prediction, to capture the DNA sequencing patterns corresponding to specific cPeak shapes (**Fig. S2e**, Methods). The network independently predicts whether each edge of a cPeak is well-positioned and achieved an AUC of 86.56% on the test set when trained on labeled cPeaks. We applied the model to cPeaks with unlabeled shapes and used the newly labeled, well-positioned cPeaks for TF enrichment analysis. The enriched TF results from this analysis strongly aligned with those obtained from previously labeled cPeaks (Spearman correlation coefficient = 0.76) **(Fig. S2f)**, demonstrating the network’s capability to classify cPeak shapes based on DNA sequence patterns. Incorporating these predictions, we classified 312,497 (18.86%) as “well-positioned”, 366,716 (22.13%) as “asymmetrically-positioned”, and 977,981 (59.01%) as “weakly-positioned”. These annotations were integrated into the cPeak resource to support further analyses and applications.

### Enhancing cell type annotation of scATAC-seq data with cPeaks

We evaluated the effectiveness of using cPeaks as a reference for annotating scATAC-seq data, comparing it against three commonly used approaches for defining chromatin accessibility features and two pre-identified accessible region sets. The first approach divides the genome into 2,000 bp bins as basic chromatin accessibility feature units^53–55^, referred to here as “genomic bins”. The second approach aggregates all sequenced single cells into pseudo-bulk samples and calls peaks from these aggregates^30,29^, referred to as “pseudo-bulk peaks”. The third approach clusters cells based on features such as pseudo-bulk peaks or cell-sorting labels, calls peaks by combining cells within each cluster, and then unites the peaks from all clusters together as the reference features^5,56^, referred to as “united peaks”. These approaches, along with two pre-defined feature sets—cDHS and CATLAS cCREs—were evaluated against cPeaks for their performance in cell annotation tasks. For each approach, we tested different levels of feature selection, including all features, the top 50,000 highly variable features (HVFs), and the top 5,000 HVFs (Methods, **Fig. 4a**).

**Fig. 4.**
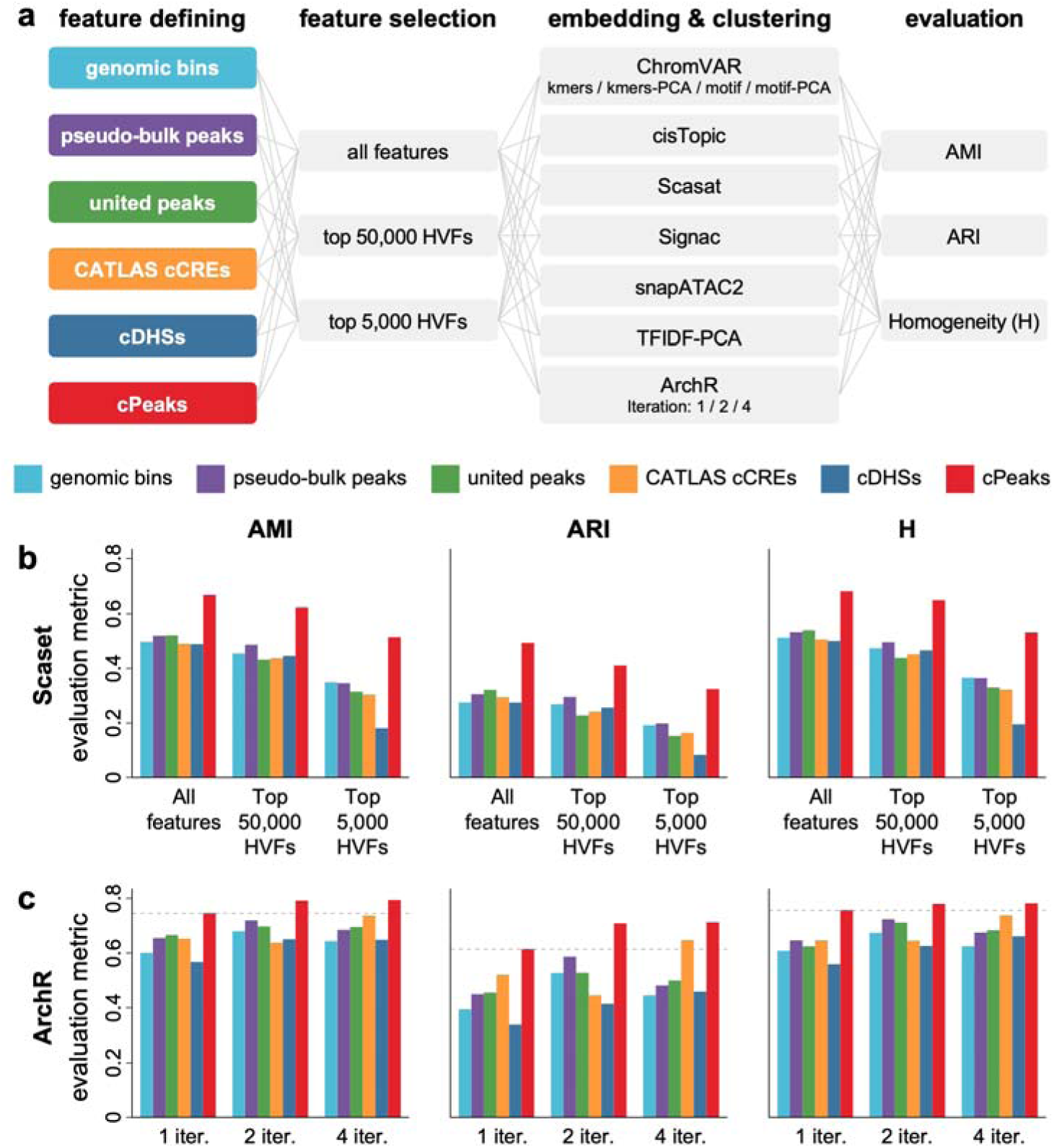
The performance of cell annotations on the FACS2-sorted hematopoietic differentiation dataset. (a) The analysis workflow of cell annotations. We experimented with six feature-defining approaches to obtain references for generating cell-by-feature matrices, three ways of feature selection, and seven strategies to embed the features before clustering analysis, and used three metrics to evaluate the results. (b) Comparison of Scaset’s performances with the six feature-defining approaches and three ways of feature selection. (c) Comparison of ArchR’s performances with the six feature-defining approaches and different iterations. Each gray dashed line represents the evaluation metric using cPeaks as a reference with one iteration.

We used an scATAC-seq dataset of hematopoietic differentiation^9^, comprising 2,034 cells with FACS-sorting labels as ground truth. For each approach, we generated cell-by-feature matrices, applied multiple embedding methods for dimensionality^19,29,41,53,57,58^, and used the Louvain method^59^ for clustering and annotation (**Fig. 4a**, Methods).

Annotation results were assessed using adjusted rand index (ARI), adjusted mutual information (AMI), and homogeneity (H)^60^ (**Fig. 4a**, Methods). Across all evaluation metrics, cPeaks consistently delivered the best or among the best performances compared to the other feature-defining approaches (**Fig. 4b and S3**). The superiority of the cPeaks became more pronounced when using the top 5,000 or top 50,000 HVFs. Considering that the definition of cPeaks was independent of this particular dataset, this performance demonstrates that cPeaks can effectively serve as a robust general reference for scATAC-seq data.

To further validate the performance of cPeaks, we employed the iterative method of ArchR^61^, which employs an iterative reduction approach to select appropriate features and refine cell embedding results. We conducted 1, 2, and 4 rounds of iteration, comparing cPeaks to the five other feature sets (**Fig. 4a**, Methods). The results showed that cPeaks outperformed other feature sets after just one iteration and continued to improve with additional iterations (**Fig. 4c**). The findings demonstrate that cPeak not only streamline the ArchR analysis workflow but also provide superior results compared to other feature sets, making cPeaks an optimal choice for cell type annotation of scATAC-seq datasets.

### Facilitating the discovery and analysis of rare cell types with cPeaks

One expected advantage of using a generic reference, such as cPeaks, rather than dataset-dependent references, is the potential to identify rare cell types, as cells of rare types often lack sufficient representation in datasets to call out their marker peaks. To evaluate this advantage, we utilized cPeaks to annotate an scATAC-seq dataset of peripheral blood mononuclear cells (PBMCs) generated by 10X Genomics (Methods). This dataset includes 8,728 cells with 87,561 pre-identified pseudo-bulk peaks. We first used cPeaks as the reference to generate the cell-by-feature matrix, and applied normalization, feature selection and dimensionality reduction following common practice (Methods).

We then performed graph-based clustering on the reduced dimension and visualized the clustering results using Uniform Manifold Approximation and Projection (UMAP) plots (**Fig. S4a**). The analysis grouped cells into multiple clusters, including some clusters containing small proportions of cells. We then transferred cell type labels from a corresponding scRNA-seq PMBC dataset through cross-modality integration using the anchor method^62^ in Seurat^63^ (**Fig. 5a**). For each cell in the scATAC-seq data, we obtained an inferred cell type with a prediction score reflecting the annotation confidence. Notably, cPeaks outperformed pseudo-bulk peaks, yielding higher prediction scores (**Fig. 5b and S4b**).

**Fig. 5.**
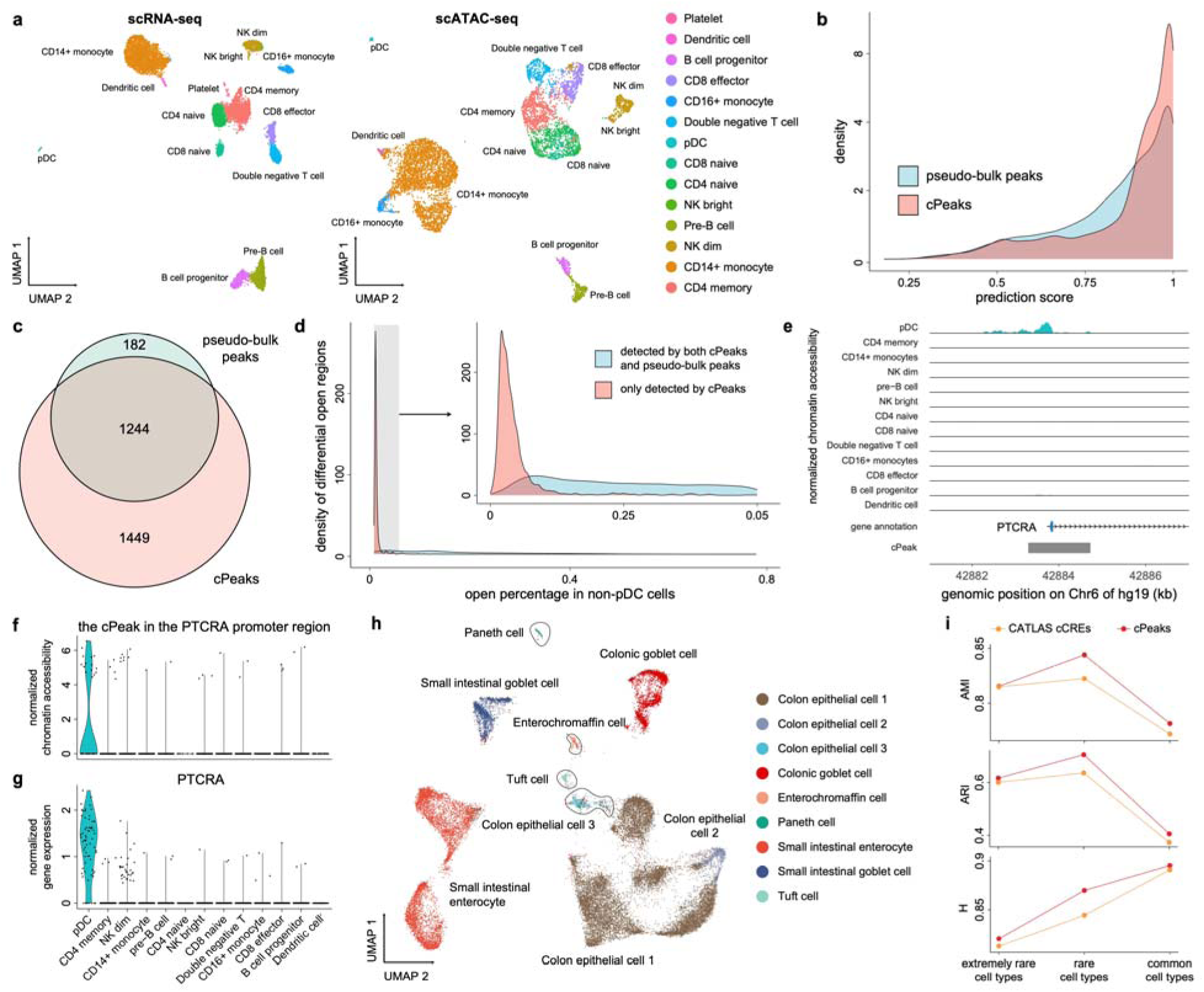
Rare cell type discoveries using cPeaks as the reference. (a) The UMAP plots of scRNA-seq (left) and scATAC-seq (right) PBMC data, with colors labeling the cell types of the cells. The scRNA-seq cells were assigned to their cell labels according to gene expression, and the labels of scATAC-seq cells were inferred using cross- modality integration. We removed cells in cluster 18 (shown in Fig. S5a) of the scATAC-seq data, as cells in the cluster showed low prediction scores on average. pDC: plasmacytoid dendritic cells; NK dim: nature killer dim cells; NK bright: nature killer bright cells. (b) Histogram of the prediction scores of the cross-modality integration between the scRNA-seq data and scATAC-seq data analyzed with cPeaks (red) or pseudo-bulk peaks (blue). The prediction score reflects the confidence of predicted labels. (c) The Venn plot of the number of differential accessible regions calculated by cPeaks (red) or pseudo-bulk peaks (blue) for pDCs. (d) The distribution of accessible percentages of two accessible region categories in non-pDC cells. The small panel on the top right is a zoom-in of the density plot of regions with less than 0.05 accessible percentage in non-pDCs. (e) The chromatin accessibility in all cell types around the promoter regions of *PTCRA*, measured by the sum of scATAC-seq reads of all cells of each type. (f) The distribution of the normalized reads aligned to the cPeak in the *PTCRA* promoter region across different cell types. The reads are normalized by the frequency-inverse document frequency (TF-IDF) method. Each dot represents a cell. (g) The normalized expression of *PTCRA* in different cell types in scRNA-seq data. The gene expression is normalized by the total expression in each cell and log-transformed. (h) The UMAP plot of subtypes in GI epithelial cells, clustered using cPeaks as the reference. The colors show the cell types provided by the original paper. The black circles mark the 4 rare cell subtypes: paneth cell, enterochromaffin cell, tuft cell and colon epithelial cell 3. (i) The performance of cell clustering on different groups of cell types, analyzed with different references.

Within the cPeaks-derived clusters, three rare cell types were identified: plasmacytoid dendritic cells (pDCs), dendritic cells, and natural killer bright (NK bright) cells, each comprising less than 1% of the total dataset (**Fig. S4c**). Taking pDCs as an example, we identified differential accessible regions specific to this cell type by comparing their chromatin accessibility with all other cells (Methods). Using cPeaks, 1,449 additional differential accessible regions were detected compared to pseudo-bulk peaks (**Fig. 5c**), These regions were specifically accessible in pDCs (**Fig. 5d and S4d**), highlighting the limitations of pseudo-bulk peaks in detecting accessible regions for rare cell types. These differential accessible regions were important for investigating the functions of rare cell types. For example, we identified the promoter region of the *PTCRA* gene as a pDC-specific accessible region (**Fig. 5e-f**). Consistently, scRNA-seq data showed that *PTCRA* expression was restricted to pDCs (**Fig. 5g**), aligning with previous studies that identified *PTCRA* as a marker of human pDCs^64,65^.

We extended this analysis to rare subtypes within cell groups, focusing on gastrointestinal (GI) epithelial cells in CATLAS^5^. They included nine subtypes, four of which represented less than 2% of the cells (**Fig. S5a**). Pseudo-bulk peaks in the original study failed to separate these subtypes (**Fig. S5b**), requiring an additional peak calling step to resolve rare subtypes (**Fig. S5c**). In contrast, replacing pseudo-bulk peaks with cPeaks as the reference enabled the distinction of all subtypes in a single step (**Fig. 5h**).

We further employed cPeaks to reconstruct the whole CATLAS which encompasses 30 adult tissue types^5^. CATLAS includes a pre-defined unified feature set, CATLAS cCREs, which was derived from pseudo-bulk peak-calling and peak integration. To evaluate cPeaks, we performed one-round cell clustering using both cPeaks and CATLAS cCREs as features, following the strategies in the original paper^5^ (Methods). We assessed the results using 30 major cell groups and 111 cell types annotated by the original paper as ground truth. Both cPeaks and CATLAS cCREs exhibited comparable performance for the 30 major groups (**Fig. S5d**).

When clustering was refined to 111 cell types, cPeaks showed slightly superior to CATLAS cCREs on all evaluations. For example, ARI increased by 0.03, and silhouette score increased by 0.02 (**Fig. S5e**). Depending on the proportion of each cell type in the whole dataset, we categorized cell types into three parts, each with 37 cell types: extremely rare cell types (average proportion 0.1%), rare cell types (average proportion 0.3%), and common cell types (average proportion 1.2%) (**Fig. 5i**, Methods). Extremely rare cell types, constituting only 4% of the total cells, were challenging to cluster distinctly even with a complete feature set, and both cPeaks and CATLAS cCREs performed similarly. Additional clustering of cell groups was necessary to resolve these cell types, as demonstrated in our earlier GI epithelial cell analysis (**Fig. 5h**), where extremely rare cell types can be distinguished after another round of clustering using cPeaks. For common cell types, which represent 82% of all cells, cPeak slightly outperformed CATLAS cCREs with an ARI increased of 0.03. For rare cell types, cPeaks provided the most notable improvements, with ARI increasing by 0.07. This advantage underscores cPeaks’ ability to retain rare cell type-specific features that may be lost during CATLAS cCREs’ peak merging and trimming processes. By offering a more comprehensive feature set, cPeaks facilitated better clustering and representation of low-abundance cell types.

Overall, the results demonstrated that cPeaks can simplify scATAC-seq data processing and facilitate the discovery and analysis of rare cell types and their specific features which may otherwise be difficult to be detected using traditional peak-calling approaches. This capability reinforces its value as a universal reference for scATAC-seq analyses, enabling robust exploration of cellular diversity.

### Investigating cell state transitions during development with cPeaks

cPeaks were defined using the data of ATAC-seq datasets of adult healthy samples. We investigated their usability and performance in analyzing developmental data. We utilized a human fetal retinal scATAC-seq dataset encompassing early (8 weeks) and late (13 weeks) stages of retina organogenesis^33^. Using cPeaks as the reference, we generated cell-by-feature matrices, applied UMAP for dimensionality reduction, and labeled cells according to the original study (Methods). For early-stage cells, three developmental trajectories were observed, leading from multipotent progenitor cells to horizontal cells, cones, and retinal ganglion cells (**Fig. 6a**). The pseudo-time values of these cells displayed a clear progression order in the UMAP (**Fig. 6b**). For later stages, we separately analyzed samples collected from the center and the periphery of the retina, and the results showed clear clusters of cell types and their transition states (**Fig. S6a-b**). Notably, the datasets used to define cPeaks excluded fetal stages and retinal samples, demonstrating the capability of cPeaks to analyze fetal development and unexplored organs. These results suggest that cPeaks constructed from adult samples encompass most accessible regions in early developmental stages.

**Fig. 6.**
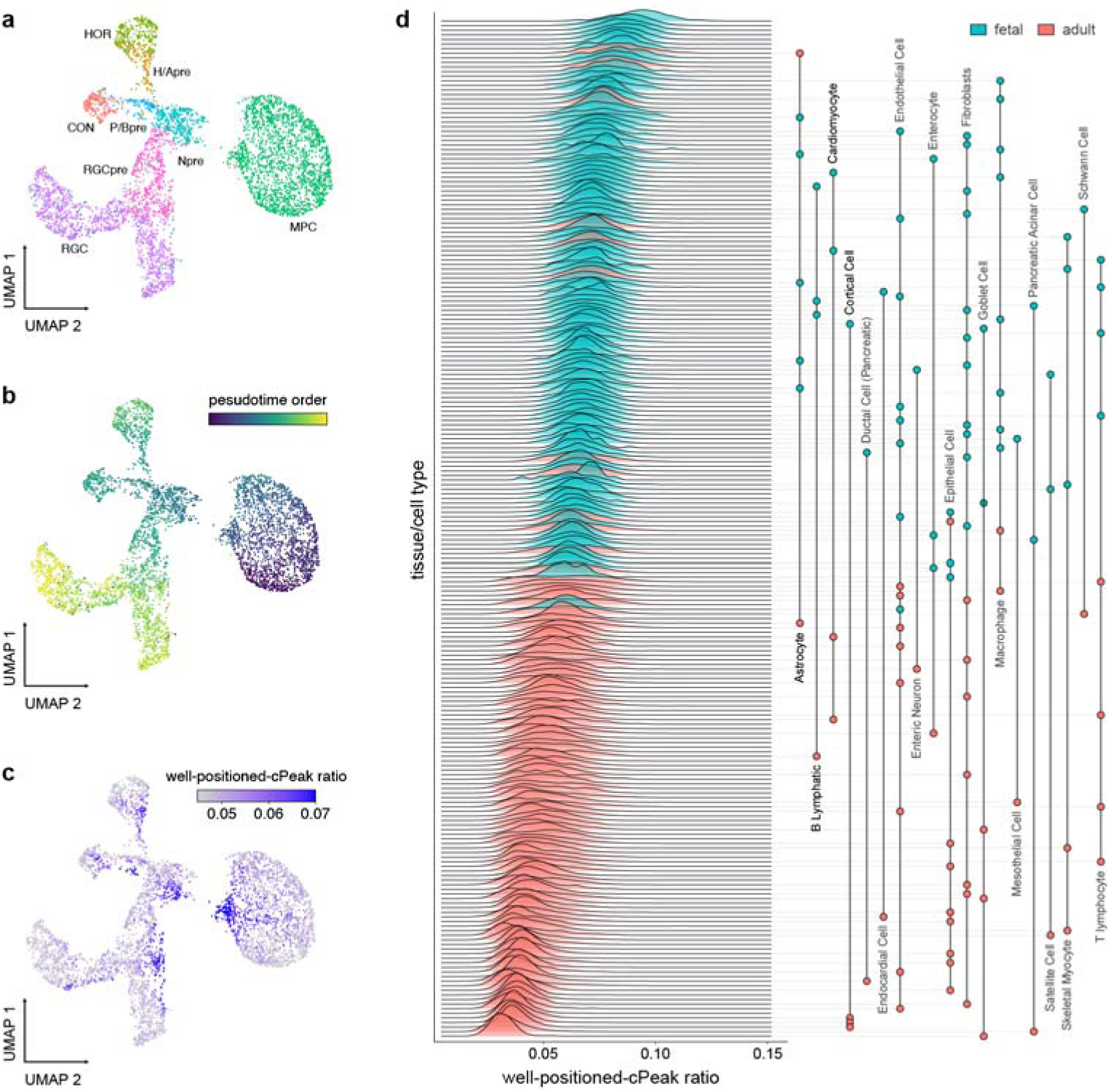
Investigating cell state transitions during development with cPeaks. (a-c) The UMAP plot of the 8-week retina scATAC-seq data colored by (a) cell types, (b) pseudotime, and (c) well- positioned-cPeak ratios. (d) Well-positioned-cPeak ratios across different cell types and life stages in scATAC-seq atlas data. Each box represents a single cell type, with colors indicating different life stages. Samples ranked by the median of well-positioned-cPeak ratios. Cell types shared in both fetal and adult samples are linked with grey lines. All the results are analyzed with cPeaks as the reference. MPC, multipotent progenitor cell; Npre, neurogenic precursor; P/B-pre, photoreceptor/bipolar precursors; CON, cones; H/Apre, the horizontal/amacrine precursors; HOR, horizontal; RGC, ganglion cells; RGCpre, RGC precursors.

We found that TFs associated with well-positioned cPeaks were enriched in GO terms associated with development, and they significant overlap with key developmental and differentiation TFs, such as pioneer factors. This observation prompted the hypothesis that well-positioned cPeaks play critical roles in cellular differentiation and development. To test this, we analyzed the accessibility dynamics of well-positioned cPeaks during development by calculating the well-positioned-cPeak ratio in all accessible cPeaks (Methods). The results revealed increased well-positioned-cPeak ratio during transition periods, followed by a decrease in accessibility at later developmental stages compared to earlier ones (**Fig. 6c and S6c-d**).

To further validate our hypothesis, we examined two large single-cell datasets: a human fetal cell atlas^66^ and CATLAS^5^ (Methods). The well-positioned-cPeak ratio was found to be higher in fetal cells compared to adult cells (**Fig. 6d**). For most cell types share by fetal and adult samples, the well-positioned-cPeak ratio consistently decreased in adult cells relative to their fetal counterparts (**Fig. 6d**). These findings underscore the association of well-positioned cPeaks with cellular differentiation and development.

### Characterizing tumor cell subtypes and tumor progression with cPeaks

We then investigated the use of cPeaks in analyzing tumor cells. We re-analyzed a human gynecologic malignancy scATAC-seq dataset of 74,621 cells in the endometrium or ovary of 11 treatment-naive cancer patients^34^, using cPeaks as the reference (Methods). The analysis demonstrated clear clustering of cells by type, with tumor cells from gastrointestinal stromal tumor (GIST), endometrial cancer, and ovarian cancer distinctly separated from normal cells (**Fig. 7a and S7a**). Labeling the cells with copy number variations (CNVs) (**Fig. 7b**) and the expression of cancer marker genes such as *KIT*, *WFDC2*, and *MUC16* (**Fig. S7b**) further validated these clusters.

**Fig. 7.**
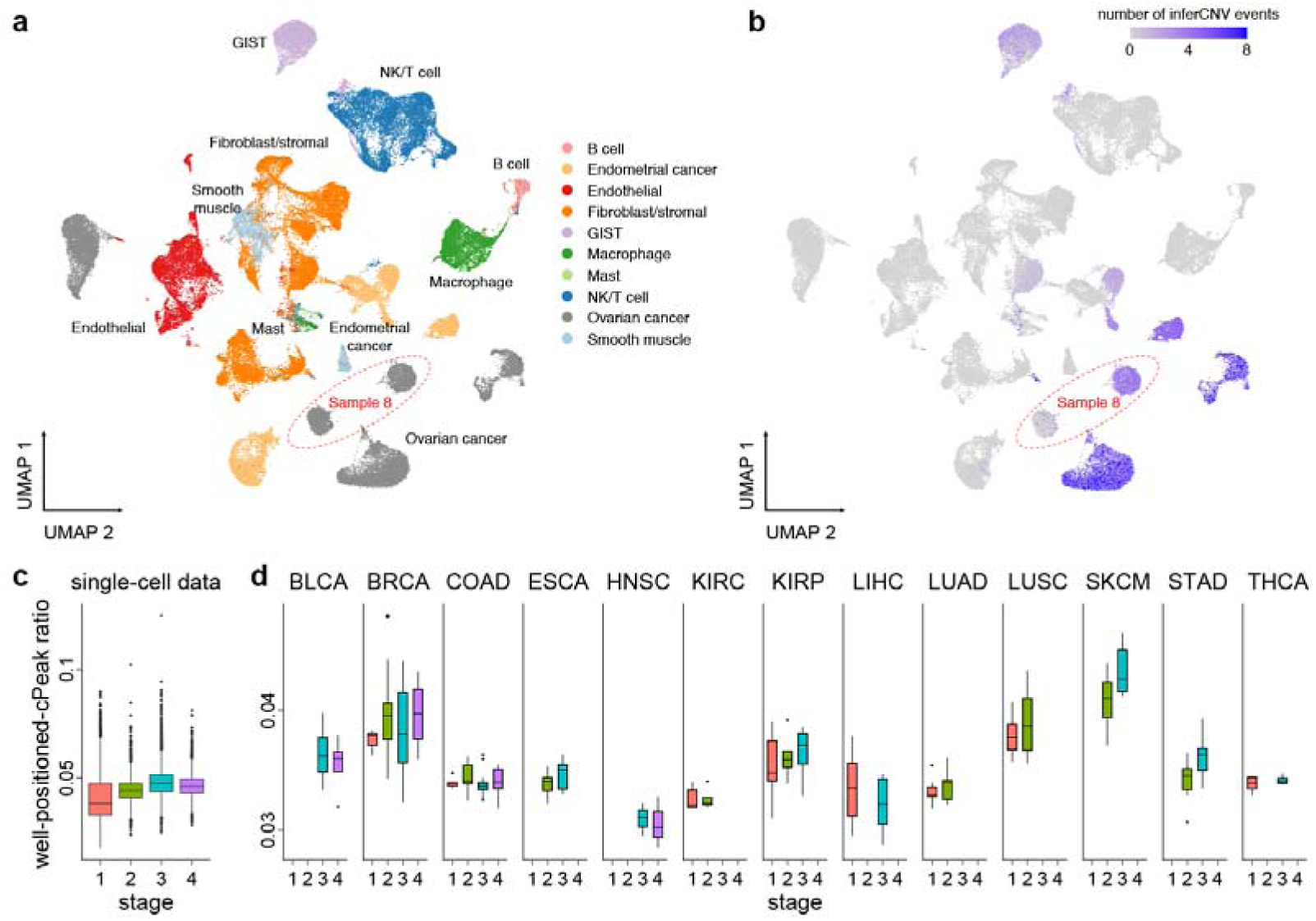
Characterizing tumor cell subtypes and tumor progression with cPeaks. (a) The UMAP plot of the human gynecologic malignancy scATAC-seq data. We aggregated all cells in the endometrium and ovary from 11 patients. Colors represent different cell types. Among them, GIST, endometrial cancer, and ovarian cancer are cancer cells. Ovarian cancer cells in sample 8 are circled by the red dashed line. GIST, gastrointestinal stromal tumor; NK/T cell, nature killer cell or T cell. (b) The UMAP plot of the human gynecologic malignancy scATAC- seq data colored by inferred CNVs. The ovarian cancer cells provided by sample 8 are circled by red dashed line. The two sub-clusters in these cells have different CNVs. All the results are analyzed with cPeaks as the reference. (c) Well-positioned-cPeak ratios in the human gynecologic malignancy scATAC-seq data across different tumor stages. (d) Well-positioned-cPeak ratios in the pan-cancer TCGA dataset across different tumor stages.

Beyond replicating the findings from the original study, we observed that ovarian cancer cells in sample 8 formed two distinct clusters (**Fig. 7a and S7a**). Both clusters expressed the cancer marker *WFDC2* (**Fig. S6b**), but only one cluster exhibited significant CNV enrichment (**Fig. 6b**). This observation suggested the presence of two subclones with distinct genomic properties within the tumor, highlighting the potential of cPeaks to identify subclusters in tumor cells. The generality of cPeaks as a reference may reveal subtle cellular differences that could be missed using dataset-derived references.

We further examined the accessibility dynamics of well-positioned cPeaks in various tumor cells. **Fig. 7c** highlights the differences in well-positioned-cPeak ratios across tumor stages revealing a trend of higher proportions of well-positioned cPeaks in later-stage tumors. To validate this, we analyzed the Pan-Cancer TCGA dataset^16^ which includes 23 tumor types and 404 datasets. Consistent with our hypothesis, most cancers exhibited increased well-positioned-cPeak ratios in later stages, with exceptions in head and neck squamous cell carcinoma (HINSC) and liver hepatocellular carcinoma (LIHC) (**Fig. 7d**).

Integrating these tumor findings with the developmental analyses, we proposed that well-positioned cPeaks may serve as a hallmark of cellular proliferation and differentiation potential. These cPeaks show high activity in fetal cells, reach their lowest accessibility in adult cells, but are reactivated and become highly accessible in tumor cells. Moreover, their accessibility proportion steadily increases during tumor progression. These results highlight the dynamic regulatory role of well-positioned cPeaks in development and disease, emphasizing their significance in cellular state transitions and tumorigenesis.

### cPeaks as a generic reference for ATAC-seq data analysis

We developed a comprehensive resource for cPeaks, including basic information and annotations (**Table S4**). The basic information includes cPeak IDs and genomic locations, while the annotations describe the housekeeping status and shape patterns. To enhance usability, we linked cPeaks with the other widly used accessible region sets and regulatory databases, namely cDHS^37^, CATLAS cCREs^5^, and ReMap^67,68^, offering relevant information about regulatory vocabularies, cell-type specificity, and binding TFs (Methods).

To facilitate the practical application of cPeaks, we developed multiple computational interfaces that integrate with existing scATAC-seq analysis platforms. Specifically, cPeaks have been incorporated into SnapATAC2^69^ and ArchR^61^, two widely used packages for single-cell epigenomics analysis. SnapATAC2, implemented in Python, supports efficient construction of cell-by-feature matrices, where cPeaks serve as a standardized reference, enabling consistent feature selection across datasets. In ArchR, an R-based platform, cPeaks can replace default peak or bin sets to generate cell-by-feature matrix. Both platforms allow for the direct integration of cPeaks into their workflows, streamlining the analysis of chromatin accessibility in single-cell datasets.

In addition to these platform-specific integrations, we provide a standalone Python script for researchers requiring additional flexibility in their workflows. This script enables the transformation of fragment files into cPeaks-based data matrices and transformation of cell-by-peak matrices into cell-by-cPeak matrices, offering a straightforward method to incorporate cPeaks into custom pipelines.

By integrating cPeaks into established analysis platforms and offering flexible standalone solutions, we provide a robust framework for analyzing scATAC-seq data. Comprehensive documentation, tutorials, and code examples for all tools are available at https://mengqiuchen.github.io/cPeaks/Tutorials.

## Discussion

scATAC-seq generates large, sparse datasets from thousands of cells, where direct peak calling on each cell is infeasible. Combining cells into pseudo-bulk samples can compromise single-cell resolution and obscure rare cell features. The scale of scATAC-seq also makes feature integration labor-intensive. A reference set can help address these challenges by reducing dependence on pseudo-bulk methods, preserving single-cell details, and providing a consistent framework for integrating data across studies.

In this study, we identified a shared set of potential chromatin accessible regions across different cell types, offering a useful resource for improving scATAC-seq analysis. Through merging the peaks from multiple bulk ATAC-seq datasets, we established the concept of consensus peaks (cPeaks), and constructed a reference set comprising 1.4 million observed cPeaks. To expand the reference to accommodate unseen cell types with unique accessible regions, we developed a deep CNN model to predict cPeaks from DNA sequence patterns and identified 0.28 million predicted cPeaks. Together, these observed and predicted cPeaks constitute the first cPeak reference for chromatin accessibility. The consistent shapes, genomic localization, and predictability of cPeaks across diverse cell types and experimental conditions suggest that cPeaks represent an inherent feature of the genome.

We conducted a series of experiments to validate the effectiveness and advantages of cPeaks as a generic reference for scATAC-seq data analyses. The results demonstrated that cPeaks perform on par with or surpass the best existing approaches in key downstream tasks such as cell type annotation. Notably, cPeaks exhibited superior sensitivity in identifying rare cell types or subtypes from scATAC-seq data. Furthermore, the cPeaks derived from healthy adult samples proved to be effective when applied to data from early developmental stages and cancers. By categorizing cPeaks according to their shapes, we identified well-positioned cPeaks may serve as hallmarks of cellular proliferation and differentiation potential. These findings suggest that systematically annotating accessible regions using cPeaks can facilitate comprehensive biological discovery.

The success of cPeaks as a generic, data-independent reference across diverse tasks underscores the existence of a shared set of accessible regions in the human genome across different cell types. We propose that future ATAC-seq or scATAC-seq data can be analyzed by first aligning the sequencing reads to cPeaks. This approach positions cPeaks as analogous to the transcriptome reference in scRNA-seq analysis, enabling standardized and uniform data comparison and integration across studies. Reads that do not align to cPeaks can be further investigated through specialized analyses to further identify unique accessible regions and their associated functions, and these regions can subsequently be incorporated into cPeaks, continuously improving its coverage and utility.

Currently, the emergence of foundation models based on transcriptions has significantly advanced our understanding of gene regulation and cell behaviors^70^. Similarly, establishing foundation models for chromatin accessibility data is crucial. A major challenge lies in the lack of standardized features across datasets. The introduction of cPeaks provides a groundwork for the development of foundation models on chromatin accessibility data and even multi-omics data.

We acknowledge that the set of cPeaks we built in this work is only an initial version based on the currently available data. There is substantial room for refinement in defining and annotating cPeaks, and in uncovering their regulatory roles. Extending the cPeaks framework to other species and studing conserved chromatin accessibility signatures could provide valuable insights into the evolution of regulatory elements over time. We envision that cPeaks will promote our understanding of chromatin accessibility and its role in gene regulation by enhancing the analysis and interpretation of scATAC-seq data.

## Methods

### cPeaks generation

#### Collecting ATAC-seq data for cPeak generation

We collected bulk ATAC-seq data from ENCODE^39^ and ATACdb^40^, focusing on datasets from healthy adult samples with more than 10,000 pre-identified peaks. For each dataset, we excluded peaks on chromosome Y (ChrY) if fewer than 20 ChrY peaks were identified, retaining the remaining peaks for further analysis. For datasets from ATACdb originally generated by hg19 reference, we used CrossMap^71^ (version 0.6.5) to convert peak positions to hg38 reference. Peaks that underwent length changes during the conversion were excluded for further analysis.

#### Defining merged peaks

Merged peaks were generated by joining overlapping peaks across datasets. To leverage the high-quality datasets from ENCODE and the broad cell type coverage of ATACdb, we generated merged peaks for ENCODE and ATACdb separately. The final merged peaks consisted of two parts: all ENCODE merged peaks, and all ATACdb merged peaks that were not overlapped with any ENCODE merged peak.

#### Defining shapes of merged peaks

For each position within a merged peak, we calculated an accessible score, defined as the normalized proportion of datasets in which the position was covered by peaks, reflecting its accessibility across datasets. These accessible scores were arranged sequentially along the 5’-to-3’ direction of the merged peak, forming a continuous profile that we referred to as the “cpeak shape”. This shape captures the distribution of accessibility across the peak, providing a detailed representation of both consistent and variable accessibility patterns across datasets. When defining shapes of merged peaks on ChrY, we accounted only for the number of datasets retaining ChrY as the denominator.

#### Evaluating the consistency of peak shapes

We evaluated the consistency of merged peak shapes across tissues, sequencing techniques, peak-calling methods, and peak-calling parameters. First, we compared merged peaks generated by 48 blood samples with merged peaks generated by 101 solid tissue samples in ENCODE to assess consistency across tissues. Second, we analyzed merged peaks generated by 149 ENCODE samples with merged peaks generated by 733 DNase-seq samples used for generating for cDHS^37^ to assess consistency across sequencing techniques, downloaded from https://www.encodeproject.org/annotations/ENCSR857UZV/. The DNase-seq samples were processed using the same procedure employed for generating merged peaks from ATAC-seq data.

Third, we compared merged peaks generated by 149 ENCODE samples with merged peaks generated by 17 10X scATAC-seq samples to assess the consistency across peak-calling methods, which used MACS2^28^ and CellRanger^29^ for peak calling, respectively. The 10X samples were downloaded from

https://www.10xgenomics.com/datasets/10k-human-pbmcs-atac-v2-chromium-controller-2-standard. Finally, we examined consistency across peak-calling parameters by comparing ENCODE merged peaks and ATACdb merged peaks, which were processed using different parameter settings. For each comparison, we selected merged peaks from ENCODE with shape maxima exceeding 0.3 and calculated the PCCs for each selected region. Spline sampling was applied to equalize the lengths of each shuffled pair before calculating the PCCs. To assess significance, we performed permutation tests by randomly shuffling the pairs of merged peaks 100 times within the selected regions.

#### Generation of cPeaks from merged peaks

We used the Hartigan’s dip test^72^ (the “dip.test” function in the diptest package (version 0.76-0) in R) to measure the multimodality of merged peak shapes. We defined merged peaks with *p*-values less than 0.1 as multimodal ones. These multimodal peaks were then split into multiple unimodal peaks based on their shapes. For each position within a multimodal peak, we calculated the number of datasets where it was accessible and created an equivalent number of tuples. The first dimension of each tuple corresponds to the position, while the second dimension is a value randomly sampled from a standard normal distribution. In this way, we transformed the shape of each peak into a point cloud in a two-dimensional space. Then, we performed Hierarchical Density-Based Spatial Clustering of Applications with Noise (HDBSCAN)^73^ to cluster the point cloud using the “hdbscan” function in the dbscan package (version v1.1-11) in R with variable minimum size of clusters (minPts) set as the number of points dividing by the merged peak length multiplying by 50. We used the cluster ID calculated by HDBSCAN to split the merged peak into several separate regions. For each separated region, we decreased the value of minPts by 5 and repeatedly performed clustering and splitting until all separated regions were unimodal or achieved the minimum length of 800 bp. After, all the separated regions were regarded as cPeaks generated from the multimodal merged peak.

### Assessing the availability of cPeaks as a chromatin accessibility reference

To analyze the distribution of sequencing reads around cPeak centers, we aligned the centers of all cPeaks and collected reads mapped within ±1000 bp of these centers to generate an aggregated distribution. The frequency of read counts at each position was calculated and normalized by dividing it by the total number of aggregated reads. For comparing cPeaks with cDHS in terms of coverage over ATAC-seq accessible regions, we downloaded DNase-seq and ATAC-seq data of GM12878 from the ChIP-Atlas website^44^. Data files were filtered by setting the “significant” parameter to 50 and restricting peak numbers to between 30,000 and 100,000. A total of 65 datasets were obtained, comprising 62 ATAC-seq datasets and 3 DNase-seq datasets. To ensure comparability, we paired each DNase-seq dataset with the ATAC-seq dataset having the closest number of peaks, resulting in three pairs: SRX5766093 & SRX069089, SRX7124440 & SRX069213, and SRX5766134 & SRX069213. Within each pair, regions uniquely identified by ATAC-seq or DNase-seq were defined as ATAC-seq- or DNase-seq-specific accessible regions, respectively.

### Expanding cPeaks through deep learning

#### Model architecture

The architecture of the deep CNN model was modified based on scBasset^45^, a one-dimensional (1D) CNN that included six 1D convolutional blocks followed by two linear blocks. Each convolutional block includes a 1D convolution layer, batch normalization, dropout, max pooling, and GELU activation function, while each linear block consists of a linear layer, batch normalization, dropout, and GELU activation function. The convolutional layers have 288, 323, 362, 406, 456, and 512 kernels, with kernel sizes of 17, 5, 5, 5, 5, and 5, respectively. The final convolutional output of 5,120 dimensions is reduced to 32 through a linear layer, and then to 2 dimensions by another linear layer, enabling binary classification.A softmax activation function is applied at the final layer to generate class probabilities.

#### Generation of samples

Observed cPeaks shorter than 100 bp were filtered out, and longer cPeaks were chunked into segments of 2,000 bp, defining the positive samples. Using the “bedtools shuffle” function from BEDTools version 2.31.1^74^, the positions of these positive samples were randomized across the human genome, avoiding blacklist regions, to create the negative samples. The “bedtools nuc” function^74^ was applied to compute the nucleotide composition of the genomic positions corresponding to both sample types, excluding any samples containing ‘N’. The remaining samples were converted into their respective base sequences for further analysis.

#### Model training

The sequences of cPeaks (positive samples) and negative samples were encoded using one-hot encoding. Their reverse-complement sequences were also included in the training process with the corresponding labels to enhance model robustness. The data was randomly split into training, validation, and test datasets in an 8:1:1 ratio. The model was trained to classify the samples, assigning positive samples a label of 1 and negative samples a label of 0. The Adam optimizer was used with a learning rate of 0.001 and a binary cross-entropy loss (BCELoss) function. Training was performed in batches of size 64. Early stopping was applied based on an increase in validation loss, and performance was evaluated on the test dataset.

#### Evaluating model’s performance of predicting missing cPeaks

For each cell type or tissue, we masked its specific cPeaks, treating the remaining cPeaks as positive samples. Negative samples were generated from background regions, including those corresponding to the masked cell type or tissue-specific cPeak regions to simulate their absence. After preparing the positive and negative samples, we trained the CNN model and used it to predict the accessibility scores of the “lost” cell type or tissue-specific cPeaks. Histograms of the accessibility scores were plotted for each cell type or tissue, and the mean elbow point of 0.872 was used as the threshold to predict additional predicted cPeaks.

#### Predicting additional cPeaks

The trained model was subsequently used to predict additional cPeaks. Candidate sequences were generated by sliding a 500 bp window with a 250 bp stride across background regions, excluding regions on chrX, chrY, or shorter than 100 bp. These candidate sequences were fed into the trained model to compute accessibility scores, and sequences with scores exceeding the threshold defined by the mean elbow point were selected as predicted cPeaks.

#### Predicting cPeak shapes

The base sequence corresponding to the genomic positions of cPeaks was used as input, with shape categories at each endpoint serving as labels. To ensure label balance, we included all well-positioned samples and randomly selected an equal number of weak-positioned ones. The CNN model used the same architecture as before, excluding the final activation function. Input sequences were one-hot encoded, and the data were split into training, validation, and test sets in an 8:1:1 ratio. The network was trained using the AdamW optimizer and BCELoss criterion, with a batch size of 128 and a learning rate of 0.001. Early stopping was employed when validation loss increased, and performance was assessed on the test set. Using the trained CNN, we predicted the shape categories of all unlabeled cPeaks. Outputs for each endpoint were plotted across the genome, and a kernel density estimate (KDE) plot was generated. A threshold was determined based on the trough between two peaks in the KDE plot. This threshold was applied to classify the shape categories of all cPeaks, including predicted cPeaks.

### Annotating cPeaks

#### Genomic annotations of cPeaks

We used the ChIPSeeker package^75^ (version 1.35.1) to annotate cPeaks with genomic annotations. we used the “plotPeakProf2” function to compare the observed cPeaks with genomic annotations of transcriptional start sites (TSSs) and transcriptional end sites (TESs). We discretized the genomic region into 800 bins and using transcript annotations (TxDb) to map the cPeaks to TSSs and TESs. Then, we defined the promoter region as ±2,000 bp around TSS and annotated cPeaks using the “annotatePeak” function. The annotation database was EnsDb.hg38.v86 provided by the ensembldb package^27^. ChIPSeeker categorized cPeaks into seven types: promoter, 5’ UTR, 3’ UTR, exon, intron, downstream (≤ 300 bp downstream of the gene end), and distal intergenic regions.

#### Inferring regulatory roles of cPeaks

we categorized cPeaks based on their overlap with genomic and epigenetic features. cPeaks overlapping any gene promoter region were defined as known promoters, while those overlapping 3’ UTRs were annotated as UTRs. To identify cPeaks associated with CTCF binding sites, we analyzed 210 CTCF ChIP-seq data from ENCODE^39^ and classified cPeaks overlapping any CTCF ChIP-seq signals as CTCF binding sites. Histone modification data from ENCODE^39^ were also incorporated, using their called peaks as signals. cPeaks overlapping H3K4Me1 signals were predicted as enhancers, while cPeaks overlapping H3K4Me3 signals but not overlapping known promoter regions were categorized as predicted promoters.

#### Annotating housekeeping cPeaks

We used a pre-defined housekeeping geneset^46^, downloaded from https://www.tau.ac.il/~elieis/HKG/HK_genes.txt. A contingency table was constructed to quantify the number of housekeeping and non-housekeeping genes associated with housekeeping and non-housekeeping cPeaks, considering only protein-coding genes. We used Fisher’s exact test to calculate the significance of the contingency table. For housekeeping cPeaks annotated as known promoters, the associated genes were defined as those driven by these promoters. Functional enrichment analysis of GO biological process terms was performed on these genes using the “enrichGO” function in the ChIPSeeker package^75^. Enriched pathways were further summarized using the “simplify” function in the clusterProfiler package^76^.

#### Motif enrichment analysis

Only the endpoints of unimodal peaks were selected for this analysis. We extracted the 200 bp sequences from the inner sides of the cPeak endpoints. We used “findMotifsGenome.pl” in Homer to perform motif enrichment analysis, with sequences derived from well-positioned edges as the experimental group and those from weakly-positioned edges as the background. We used “annotation.pl” in Homer to annotate the enriched motifs^77^. A KDE plot was generated using seaborn.kdeplot^78^ to visualize the distribution of motifs around the endpoints. We filtered TFs associated with the enriched motifs with an adjusted *p*-value of less than 0.1 for domain and GO enrichment analysis. We used the enrichR library^79^ for enrichment analysis. For domain enrichment, we used the GO_Biological_Process_2023 database^80^ and applied an adjusted *p*-value threshold of 1e–5.

Subsequently, we clustered the enriched GO terms using the “GO_similarity” function in the “simplifyEnrichment” library^81^ to group similar terms.

#### Enrichment of PTFs

We first gathered 32 known PTFs from the literature^50^, including FOXA1, FOXA2, FOXA3, GATA1, GATA2, GATA3, GATA4, CEPBA, CEBPB, ESRRB, POU5F1, KLF4, NEUROD1, TP53, FOS, FOSL1, FOSL2, MAFG, MAFF, MAFK, JUN, JUNB, JUND, ATF2, ATF3, ATF4, ATF6, ATF7, SPI1, ASCL1, HNF1A, and PU.1. We then cross-referenced the 440 TFs in the Homer database^82^ against this list and identified 25 PTFs. Among these, 22 were significantly enriched (adjusted *p*-value < 0.1). For the remaining TFs that were not classified as PTFs, 269 displayed significant enrichment. Fisher’s exact test yielded a *p*-value of 0.016, demonstrating a significant association between the PTFs and the well-positioned-associated TFs.

#### *de novo* motif enrichment analysis

We extended the regions ±200 bp around the endpoints of well-positioned edges in unimodal cPeaks to create the experimental group. Using Homer’s “findMotifsGenome.pl” function^82^, we performed *de novo* motif discovery on the experimental group, with randomly sampled regions serving as the background. The *de novo* motifs were then scanned across the endpoints of the unimodal cPeaks using Homer’s “annotation.pl” function to annotate their occurrences.

### Evaluating the cell type annotation performance of cPeaks

#### Data preprocessing

We obtained BAM files for the FACS2-sorting hematological differentiation scATAC-seq data from https://github.com/pinellolab/scATAC-benchmarking/tree/master/Real_Data/Buenrostro_2018/input/sc-bams_nodup, which was provided by a benchmark study^60^. The dataset included 2,034 cells with the hg19 reference genome.

We generated cell-by-feature matrices using six feature sets: genomic bins, pseudo-bulk peaks, united peaks, cPeaks, cDHSs, and CATLAS cCREs. Each feature set provided genomic regions defined as accessible regions. For genomic bins, the hg19 genome was divided into fixed-length 2,000 bp bins, each treated as a feature. For pseudo- bulk peaks, we used the pre-defined peaks of the original study^9^ from the Gene Expression Omnibus (GEO) database under accession code GSE96769, where peaks were called on all aggregated cells using MACS2^28^. United peaks were derived from the peak set provided by the benchmark study^60^ from https://github.com/pinellolab/scATAC-benchmarking/blob/master/Real_Data/Buenrostro_2018/input/combined.sorted.merged.bed. For cPeaks, we excluded housekeeping cPeaks and extra-long cPeaks (> 2,000 bp) and converted the cPeaks from hg38 to hg19 using CrossMap^71^. For cDHSs, we downloaded the dataset from https://zenodo.org/records/3838751/files/DHS_Index_and_Vocabulary_hg19_WM20190703.txt.gz?download=1, which is provided in hg19 format. For CATLAS cCREs, we retrieved the hg38 version from http://catlas.org/catlas_downloads/humantissues/cCRE_hg38.tsv.gz and converted it to hg19 using CrossMap.

For each cell, sequencing reads in the BAM file were mapped to the corresponding feature set, and the number of reads overlapping each feature was counted as chromatin accessibility. The chromatin accessibility values across all features and cells were compiled into a cell-by-feature matrix for downstream analyses.

#### Selecting HVFs

We first applied TF-IDF transformation to normalize the cell-by-feature matrix. We then calculated the variance of each feature across all cells after normalization. We sorted the features by their variance and selected the top features as HVFs.

#### Embedding methods for cell representations

We utilized various embedding methods to process cell-by-feature matrices. ForChromVAR-motif^41^, we used the ChromVAR package (version 1.16.0) in R, adjusted for GC bias, resized features to 500 bp, and matched motifs from the JASPAR database^83^ using the “matchMotifs” function. The resulting cell-by-motif matrix was transformed into a cell-by-deviation Z-score matrix via “computeDeviations” in ChromVAR. ChromVAR-motif-PCA extended this method by performing principal component analysis (PCA) on the deviation matrix and selecting the top 10 principal components (PCs) for the final embedding. For ChromVAR- kmer, instead of motifs, we used 6-mers as features with the “matchKmers” function in ChromVAR, generating a cell-by-kmer deviation Z-score matrix. ChromVAR-kmer-PCA followed the same steps but added PCA to select the top 10 PCs. cisTopic^57^ (version 0.3.0) in R applied latent Dirichlet allocation (LDA) on the cell-by-feature matrix using the “runCGSModels” function, generating models with varying topic numbers as 10, 20, 25, 30, 35, and 40. The best model was selected using “selectModel”, and the final cell-by-topic probability matrix was normalized via “modelMatSelection” with “method = probability”. Scasat^58^ computed cell-cell Jaccard distances from the cell-by-feature matrix and reduced dimensions to 15 via multidimensional scaling (MDS). Signac^84^ used Seurat (v4.3.0)^63^ to create a cell-by-feature matrix, applied TF-IDF transformation, and performed latent semantic indexing (LSI) via SVD, retaining PCs 2∼50 for reduced dimensions. SnapATAC2^69^ generated a cell-by-cPeaks matrix using “snapatac2.pp.make_peak_matrix” and reduced dimensions with “snapatac2.tl.spectral”. Neighbor graphs and cell clusters were computed using “snapatac2.pp.knn” and “snapatac2.tl.leiden”, respectively, yielding 10 cell types. TF-IDF-PCA^19,53,85^ filtered sex chromosome features, applied TF-IDF transformation, and performed SVD to select the top 150 PCs as embedding features. ArchR^61^ used cPeaks as features, reduced dimensions via “addIterativeLSI”, and identified 10 cell types using “addClusters”, with iterations set to 1, 2, and 4 for comparison.

#### Evaluation

After embedding, we performed Louvain clustering^59^ with the number of clusters set equal to the number of FACS-sorted labels. The clustering results were evaluated using three metrics: Adjusted Mutual Information (AMI), Adjusted Rand Index (ARI), and Homogeneity (H). All these indexes were calculated using the “sklearn.metrics.cluster” library in Python.

### Analysis of the PBMC dataset for rare cell type discovery and analysis

We download the fragment file of the PBMC dataset from the 10x Genomic website (https://cf.10xgenomics.com/samples/cell-atac/1.0.1/atac_v1_pbmc_10k/atac_v1_pbmc_10k_fragments.tsv.gz). We mapped all fragments to cPeaks, counting the number of fragments that overlapped with each cPeak as its accessibility. This process generated a cell-by-feature matrix for cPeaks. For comparison, we downloaded the matrix using pseudo-bulk peaks as features from the same website (https://cf.10xgenomics.com/samples/cell-atac/1.0.1/atac_v1_pbmc_10k/atac_v1_pbmc_10k_filtered_peak_bc_matrix.h5). Then, we followed the Signac^84^ PBMC vignette (https://stuartlab.org/signac/articles/pbmc_vignette.html) to preprocess these two cell-by-feature matrices separately. Low-dimensional clustering and UMAP visualization were performed using dimensions 2∼30.

We annotated cells in the scATAC-seq data by integrating them with a labeled PBMC scRNA-seq dataset provided by 10x Genomics. The raw scRNA-seq data was downloaded from https://support.10xgenomics.com/single-cell-gene-expression/datasets/3.0.0/pbmc_10k_v3, and the pre-processed scRNA-seq Seurat object was downloaded from https://signac-objects.s3.amazonaws.com/pbmc_10k_v3.rds. We utilized the “FindTransferAnchors” function in Seurat^63^ for cross-modality integration and used the “TransferData” function in Seurat to infer cell type labels in the scATAC-seq dataset along with their corresponding prediction scores. After label transformation, cluster 18 was identified as having low prediction scores, with an average below 0.5. This indicated that the cells in this cluster were likely of low quality. Consequently, this cluster was excluded from subsequent differential accessibility analyses.

We performed differential accessibility analysis to find differentially accessible regions in pDCs compared to other cell types. We used “FindMarkers” in the Signac package^84^ with test.use = “LR” and latent.vars = “peak_region_fragments” to perform differential analysis.

### Analysis of the CATLAS dataset for rare cell type discovery and analysis

The fragment file and metadata of CATLAS data were downloaded from http://catlas.org/catlas_downloads/humantissues/. We first analyzed GI epithelial cells from CATLAS^5^, constructing cell-by-feature matrices with cPeaks, CATLAS-defined cCREs , and MACS2^28^, separately. For cPeaks, we used non-housekeeping cPeaks accessible in more than 50 cells as features. Peaks for GI epithelial cells were called using MACS2 with the parameters --shift 0 --nomodel --call-summits --nolambda --keep-dup all. Each matrix was preprocessed using the same steps applied to PBMCs, followed by graph-based clustering and UMAP visualization across dimensions 2 to 30.

For the analysis of the whole CATLAS, we imported 92 fragment files from CATLAS into snapATAC2^69^. Using the barcodes in the metadata download from http://catlas.org/catlas_downloads/humantissues/fragment/, we got 615,998 cells. Batch correction was performed using the “snapatac2.pp.mnc_correct” function. We used the “snapatac2.pp.select_features” function to select 500,000 features. We reduced the dimensions to 30 through the “snapatac2.tl.spectral” function and generated a neighbor graph through “snapatac2.pp.knn”. UMAP visualization was carried out using the “snapatac2.tl.umap” function. Adjusting the resolution parameter in “snapatac2.tl.leiden”, we identified either 30 or 111 cell clusters.

We calculated the proportion of each of the 111 cell types annotated in the CATLAS^5^ paper. Depending on the ratio, we categorized cell types into three parts, each with 37 cell types. Based on the ranking of cell proportions, we evenly divided these cell types into three groups, with 37 cell types in each group. Using the 111 cell type labels as ground truth, we calculated AMI, ARI, and H for each group to evaluate clustering performance.

### Analysis of the development datasets

We retrieved the fragment files and metadata for the retinal fetal dataset^33^ from GEO (accession code GSE184386) and processed the data following the filtering steps outlined in the original study. The generation of cell-by-feature matrices, clustering, and visualizations were conducted using the same methodology as applied to the PBMC dataset analysis. Cell labeling was based on the provided metadata, and all analyses were performed independently for each sample.

To quantify the well-positioned-cPeak ratio in all accessible cPeaks, we first binarized the cell-by-feature matrices, and then computed the normalized accessibility of all well-positioned cPeaks. The normalized accessibility is defined as the number of well-positioned cPeaks that are accessible normalized by the number of all accessible cPeaks.

We download the human fetal cell atlas data^66^ from http://catlas.org/catlas_downloads/humantissues/fragment/. The analysis of these data followed the same procedure as that of CATLAS data.

### Analysis of the tumor datasets

We downloaded the fragment files and metadata for the gynecologic malignancy dataset^34^ from GEO (accession code GSE173682). Cells from all eight patients were combined to generate a unified cell-by-feature matrix. We followed the preprocessing steps in the original study to filter cells. The generation of the cell-by-feature matrix, clustering, and visualizations were performed using the same methodology as applied to the PBMC dataset analysis. We labeled each cell using labels from the metadata file. The calculation of the well-positioned-cPeak ratio was performed using the same methodology as applied to the retinal fetal dataset.

For the analysis of Pan-Cancer TCGA data, we download normalized ATAC-seq sample-by-peak data as well as the stage information from https://gdc.cancer.gov/access-data/gdc-data-portal. The corresponding tumor cohort information was downloaded from https://www.science.org/doi/suppl/10.1126/science.aav1898/suppl_file/aav1898_data_s1.xlsx. Within each cohort, stages with fewer than three samples were excluded. We then transformed the sample-by-peak data into sample-by- cPeak data. Cohorts with fewer than two stages were subsequently filtered out. The calculation of the well-positioned-cPeak ratio was performed using the same methodology as applied to the retinal fetal dataset.

### The resources for cPeaks

The resource for cPeaks include basic information and cPeak annotation, organized into 15 columns:

- ID: A unique 14-character string representing each cPeak, starting with “CP”, followed by “HS” (human), and ending with a nine-digit number. For example, the first cPeak is encoded as “CPHS000000001”.
- source: Indicates the origin of the cPeak, either “observed” or “predicted”.
- chr_hg38: The chromosome where this cPeak locates in the hg38 reference genome.
- start_hg38: The start position of this cPeak in the hg38 reference genome.
- end_hg38: The end position of this cPeak in the hg38 reference genome.
- housekeeping: Specifies whether the cPeak is accessible across nearly all datasets (“TRUE” or “FALSE”).
- shape_pattern: The shape pattern of the cPeak, categorized as “well-positioned”, “asymmetrically-positioned”, or “weakly-positioned”.
- inferredElements: The inferred regulatory elements associated with the cPeak, such as “CTCF”, “TES”, “TSS”, “Enhancer” or “Promoter”.
- chr_hg19: The chromosome where this cPeak locates in the hg19 reference genome.
- start_hg19: The start position of this cPeak in the hg19 reference genome.
- end_hg19: The end position of this cPeak in the hg19 reference genome.
- cDHS_ID: The ID of the overlapped cDHS region in the hg38 reference, formatted as “chr_start_end”.
- CATLAS_ID: The ID of the overlapped CATLAS cCREs region in the hg38 reference, formatted as “chr_start_end”.
- ReMap_ID: The ID of the overlapped ReMap region in the hg38 reference, formatted as “chr_start_end”.

## Data availability

We only used public datasets in this study. The information of bulk ATAC-seq data for cPeak generation can be found in **Table S1**. The scATAC-seq and ATAC-seq data for cPeaks evaluation can be found in **Table S2**. The data used for cPeak annotation can be found in **Table S3**. Download information of all other data used for analysis was provided in Methods. The information of all cPeaks as the resource can be found in **Table S4** and can be downloaded from https://mengqiuchen.github.io/cPeaks/Tutorials.

## Code availability

All code is available on GitHub (https://github.com/MengQiuchen/cPeaks), including the generation of observed cPeaks, training of the deep CNN model, all applications on scATAC-seq data, and mapping and conversion tools for analyzing data using cPeaks. We provided a detailed tutorial on how to use cPeaks (https://mengqiuchen.github.io/cPeaks/Tutorials).

## Acknowledgments

The work is supported in part of National Natural Science Foundation of China (grants 62250005, 62373210, 62433001, 92470105), the National Key R&D Program of China (grant 2021YFF1200900), and the Tsinghua-Fuzhou Institute for Data Technology. We acknowledge the helpful discussions of Drs. Wei Xie, Xun Lan and Yinqing Li.

## Author contributions

X.Z., Q.M. and L.W. conceived the study. X.Z. and L.W. supervised the study. Q.M. and X.Wu. collected the data, designed the experiments and performed the analysis. W.C. contributed to the mapping tools and tutorial website. Y.Z. contributed to the deep CNN network. W.C., Y. Z., and C.L. tested and improved the mapping tools. Z.W., J.L. (first) and X. Wang assisted in designing the deep CNN network. X.X., Si.C., Catherine.Z., Sh.C., J.L. (second), X.L., W.X. and R.J. contributed to the interpretation of results. Q.M., X.Wu. L.W. and X.Z. wrote the manuscript. All authors read and approved the final manuscript.

