## Supplemental Figures for "A generic reference defined by consensus peaks for scATAC-seq data analysis"

### Supplementary Figures

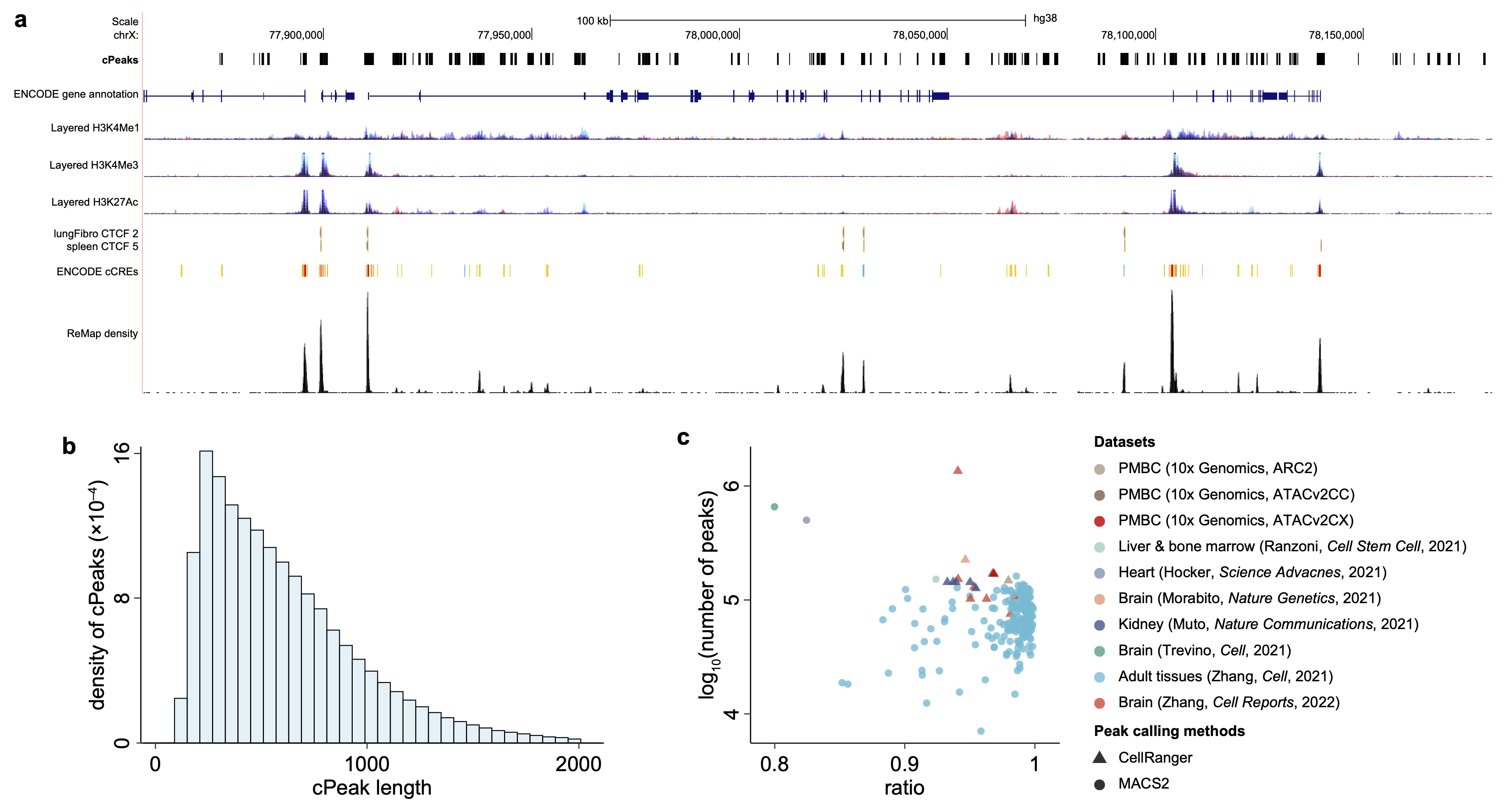

**Fig. S1. Details of observed cPeaks**. (a) An example of cPeaks in the UCSC genome browser on the regions around the PGK1 gene (chrX:77,856,686-78,181,002). The selected tracks from top to bottom were cPeaks, ENCODE gene annotation, layered H3K4Me1 marker, layered H3K4Me3 marker, layered H3K27Ac marker, lung fibroblast CTCF signal, spleen CTCF signal, ENCODE cCREs and ReMap density. The layered H3K4Me1/H3K3Me3/H3K27Ac marker combined the corresponding histone modification signals of 7 cell lines (GM12878, H1-hESC, HSMM, HUVEC, K562, NHEK, NHLF) in ENCODE together. Different colors and layers represented different cell lines. The ReMap density track represented the density of overlapped transcriptional regulators provided by ReMap Atlas. (b) The length distribution of observed cPeaks. cPeaks with lengths less than 2,000 bp were shown. (c) The coverage of observed cPeaks in 231 scATAC-seq data. The colors represented different dataset sources.

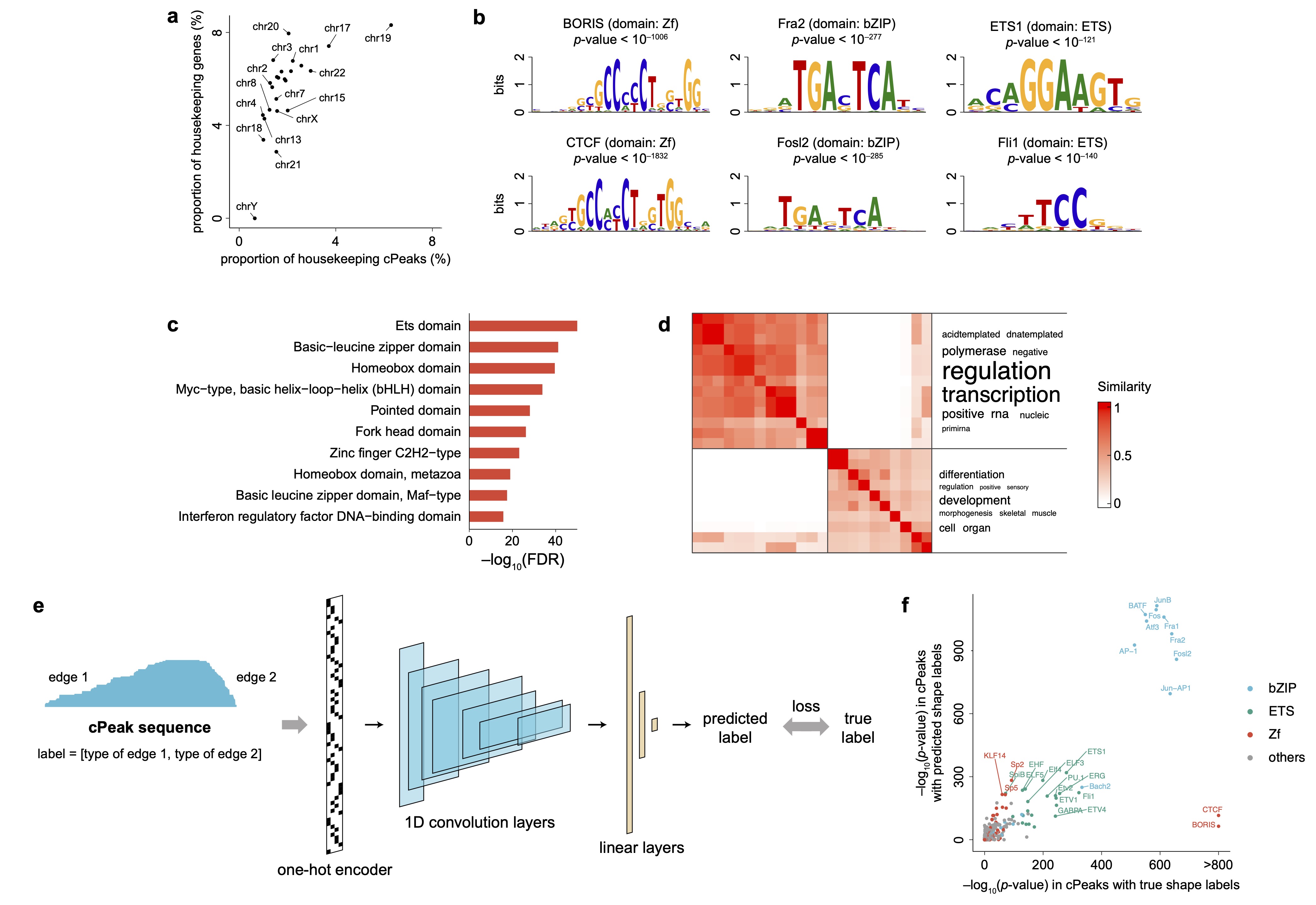

**Fig. S2. Properties of cPeaks.** (a) The proportion of housekeeping cPeaks and housekeeping genes on each chromosome. Each dot represented a chromosome. For each chromosome, the proportion was calculated by the number of housekeeping cPeaks/genes divided by the total number of cPeaks/genes on the chromosome. The PCC between the proportion of housekeeping cPeaks and housekeeping genes is 0.64. (c) Domains enriched around the well-positioned-associated TFs. The top 10 domains with the most significant adjusted *p*-values were shown. (d) The heatmap of top enriched GO terms calculated by all well-positioned-associated TFs. The row and column of heatmap represents the enriched GO terms, while the value of heatmap represents term-term similarity. (e) The CNN model for predicting cPeak shapes. (f) The correlation between the log-transformed *p*-values of enriched TFs calculated by well-positioned cPeaks with true shape labels and those with predicted shape labels. Each dot represents a single TF, with colors indicating different DNA-binding domains.

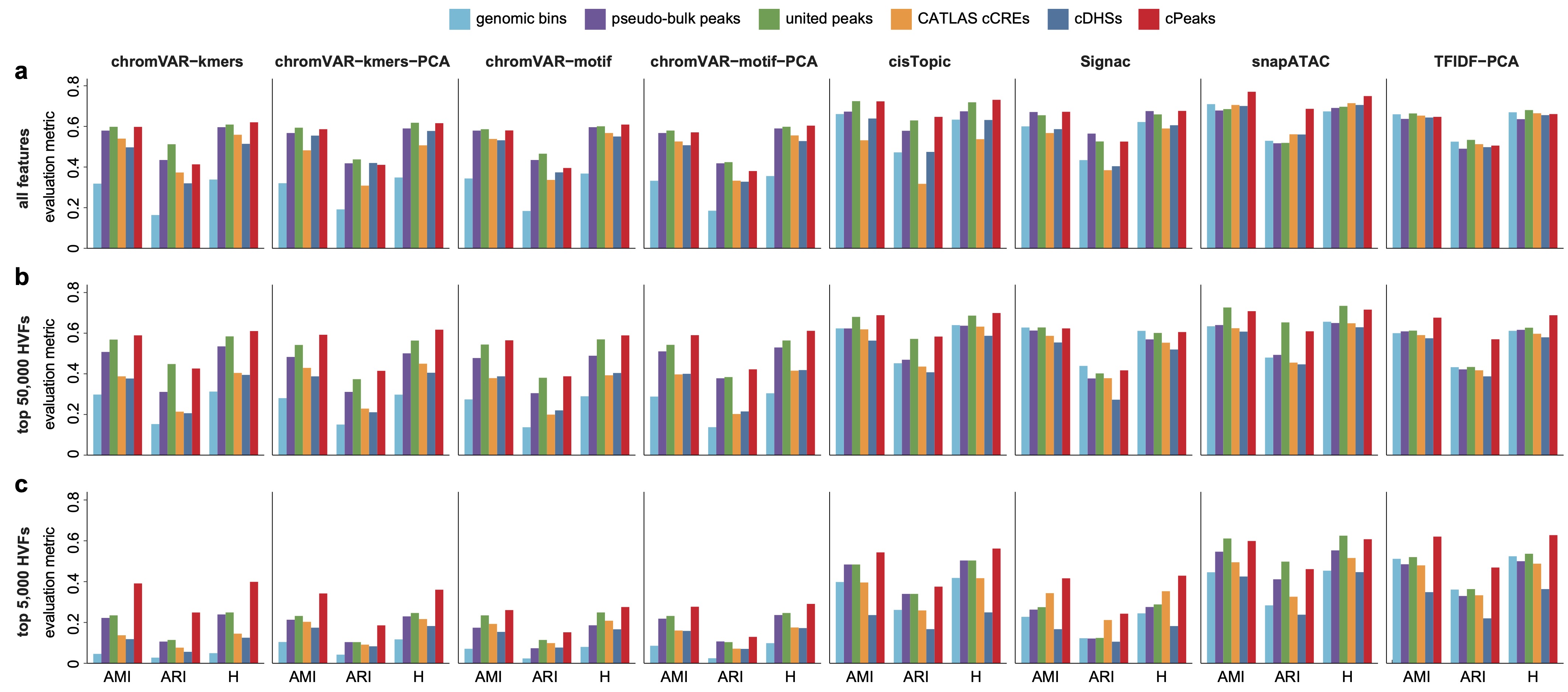

**Fig. S3. The performance of cell annotations on FACS2-sorted hematopoietic differentiation data with different evaluation metrics via different embedding methods.** (a) All features were used for analysis. (b) The top 50,000 HVFs were used for analysis. (c) The top 5,000 HVFs were used for analysis.

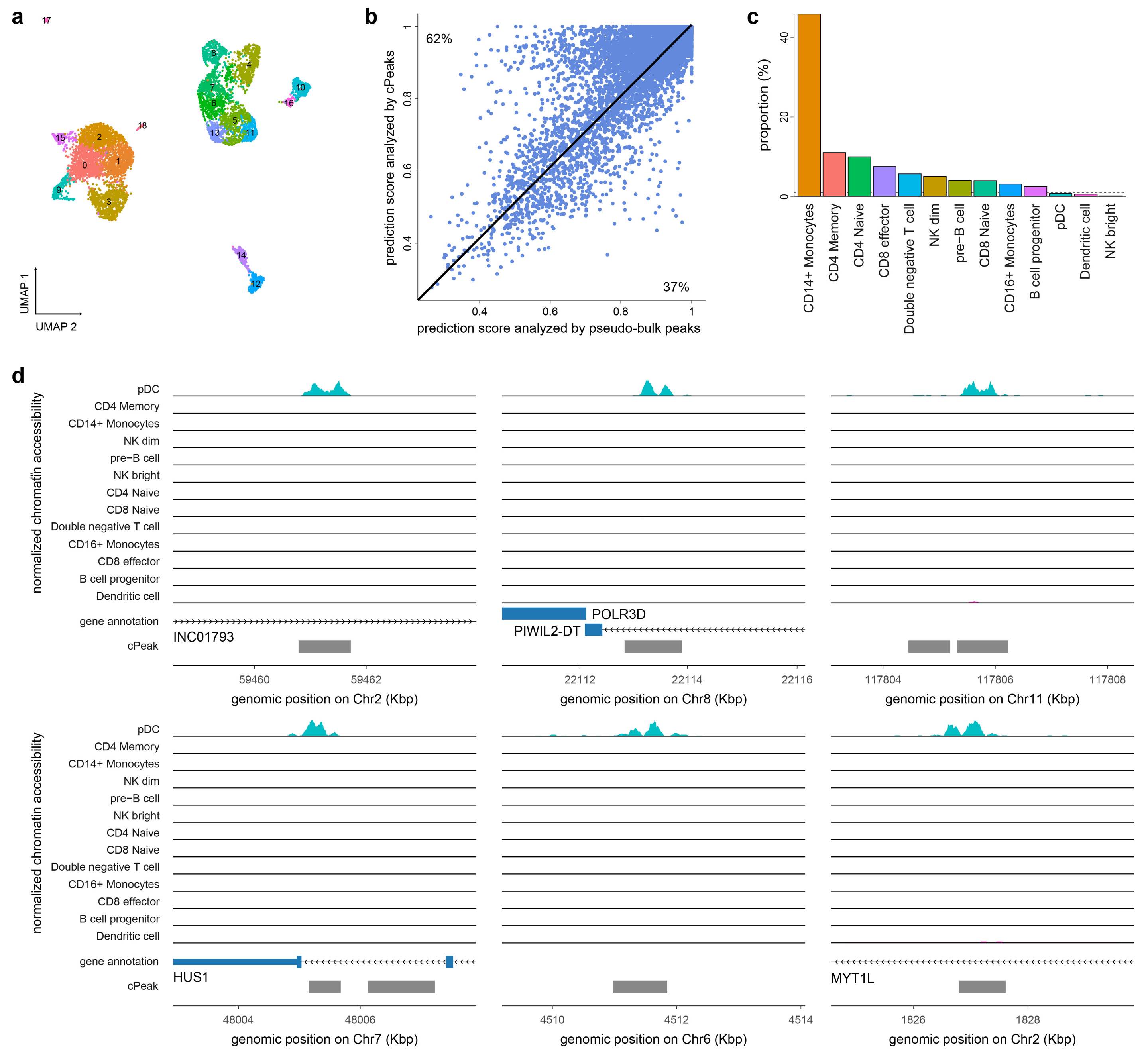

**Fig. S4. Rare cell type discovery using cPeaks on the PBMC data.** (a) The UMAP plot of the PBMC scATAC-seq data analyzed by cPeaks. Each dot represented one cell. Different colors represented different clusters. (b) Prediction scores of cells in the PBMC scATAC-seq data calculated by cPeaks and pseudo-bulk peaks. Each dot represented one cell. (c) The proportion of each cell type in the PBMC scATAC-seq data. Three rare cell types were less than 1% (the dashed black line): pDCs, dendritic cells and NK bright cells. (d) Genomic tracks of six differentially accessible regions only found by cPeaks, shown in the hg19 reference genome. Genes were annotated by RefSeq.

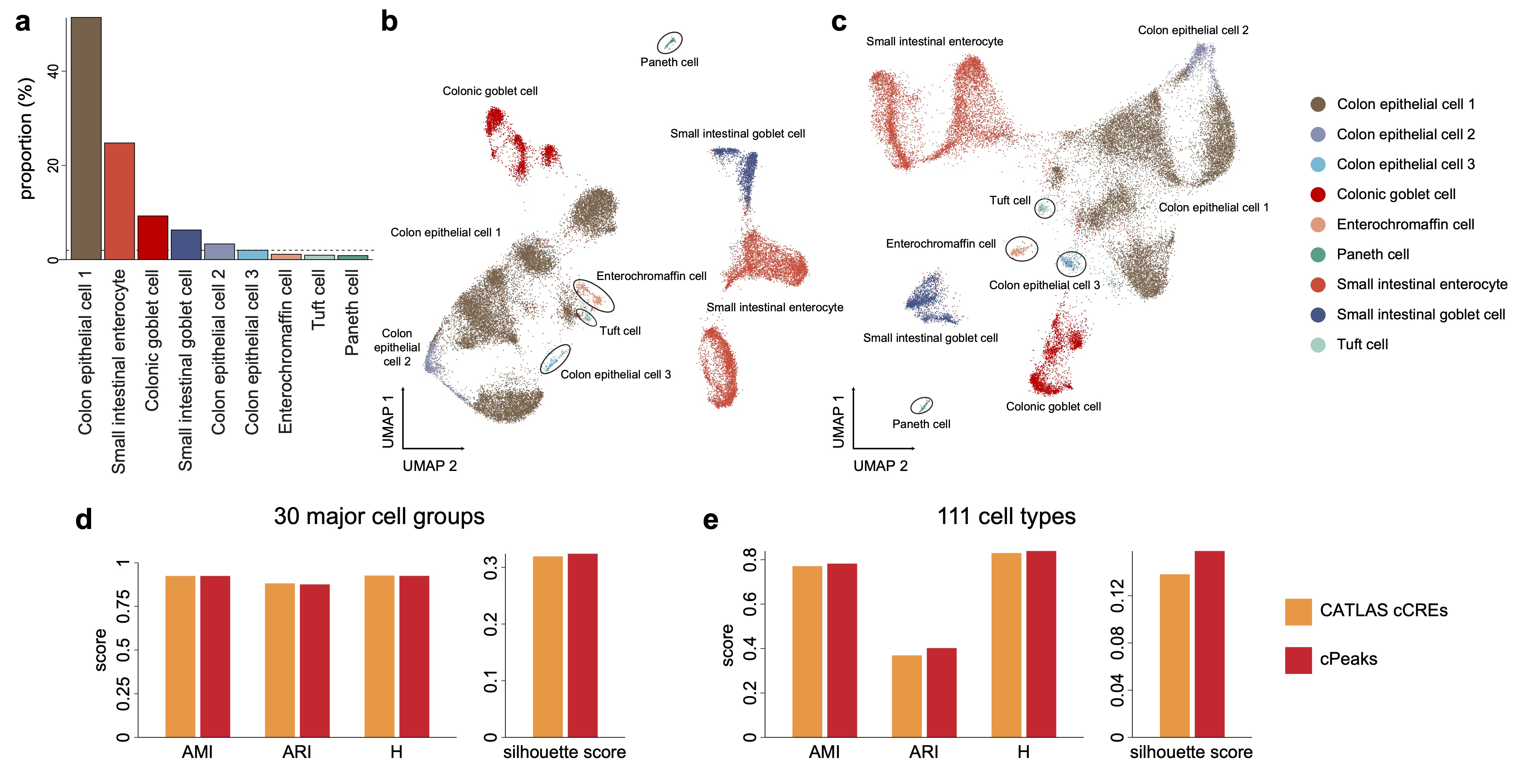

**Fig. S5. Rare cell type discovery using cPeaks on the CATLAS data.** (a) The proportion of each subtype in GI epithelial cells. Four rare subtypes were less than 2% (the dashed black line): paneth cell, enterochromaffin cell, tuft cell and colon epithelial cell 3. (b) The UMAP plot of subtypes in GI epithelial cells using merged peaks. Each dot represented one cell. Each color represented a cell type provided by the original study^5^. The black lines labeled the four rare subtypes. (c) The UMAP plot of subtypes in GI epithelial cells using pseudo-bulk peaks called by the population of all GI epithelial cells. Each dot represented one cell. Each color represented a cell type provided by the original paper. The black lines labeled the four rare subtypes. (d) Performance of cell clustering with 30 major cell groups as labels. (e) Performance of cell clustering with 111 cell types as labels.

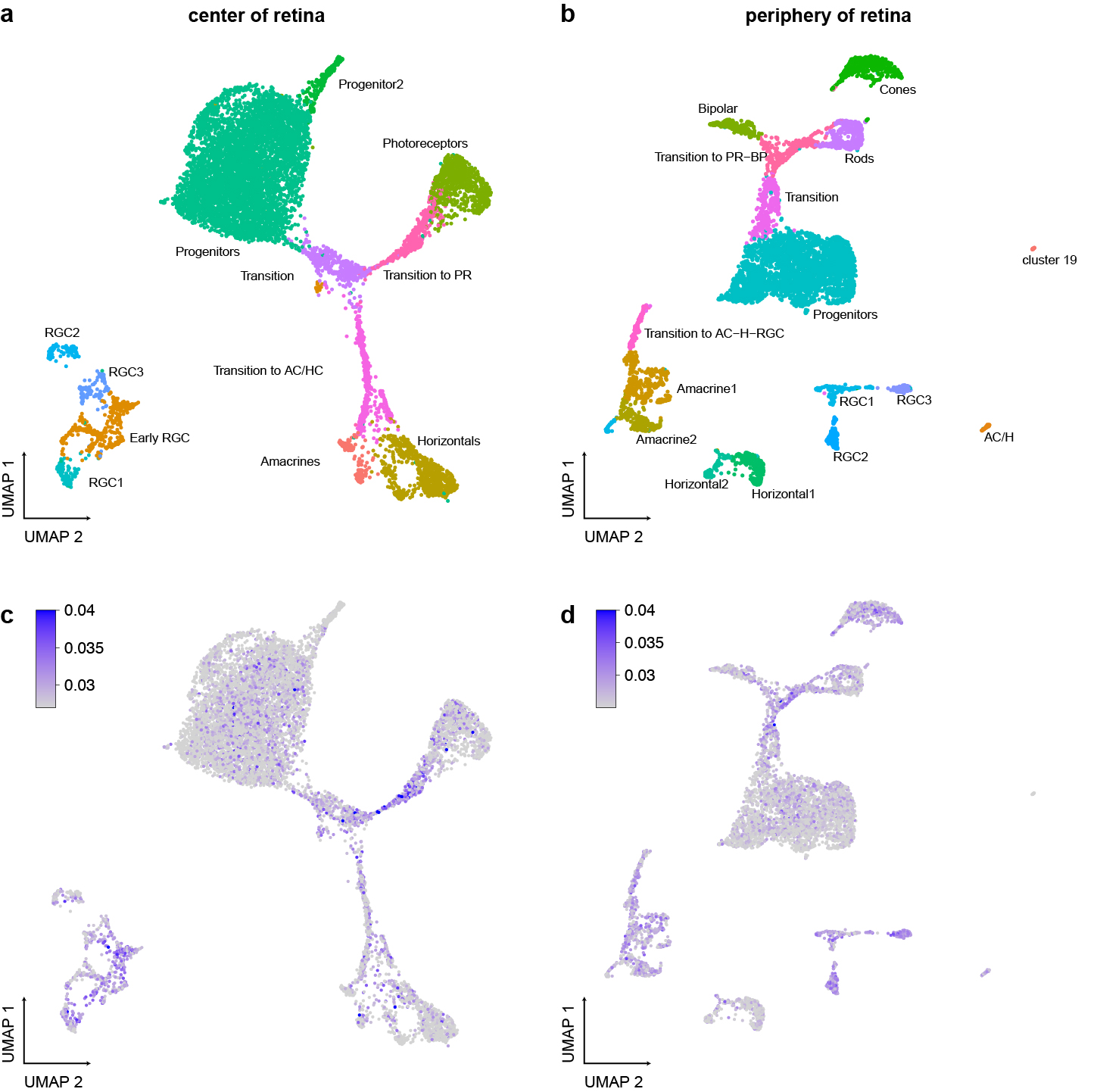

**Fig. S6. Investigating cell state transitions during development with cPeaks.**

The UMAP plot of the 13-week scATAC-seq data collected from the center of retina (a and c) and the periphery of retina (b and d). Cells are colored by cell types in (a) and (b), and by well-positioned-cPeak ratios in (c) and (d). PR, [photoreceptors](https://www.sciencedirect.com/topics/immunology-and-microbiology/photoreceptor); AC, amacrine cell; HC, horizontal cell; RGC, ganglion cell; BP, bipolar and rods; A-H-RGC, amacrine cell, horizontal cell or ganglion cell.

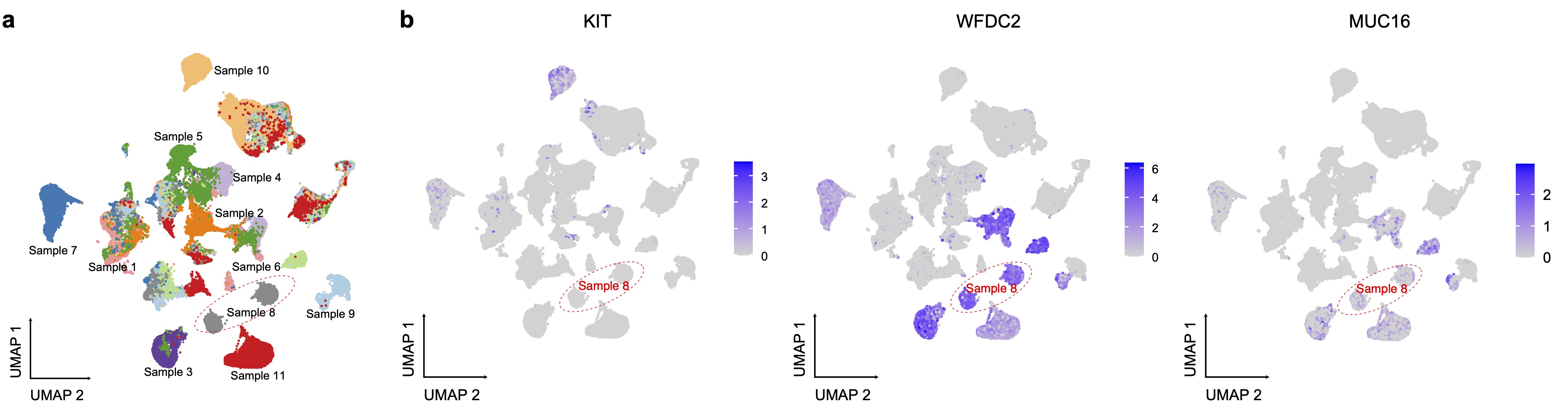

**Fig. S7. Characterizing tumor cell subtypes and tumor progression with cPeaks.** The UMAP of the human gynecologic malignancy scATAC-seq data, colored by (a) sample IDs, and (b) inferred gene expression.

### Supplementary Tables

Table S1. Information of bulk ATAC-seq data for cPeak generation.

Table S2. Information of scATAC-seq and bulk ATAC-seq data for cPeak evaluation.

Table S3. List of epigenetic and CTCF ChIP-seq data for cPeak annotations.

Table S4. Information of all cPeaks.
